# *Trans*-regulation of heterochromatin underlies genetic variation in 3D genome contacts

**DOI:** 10.64898/2025.12.10.693515

**Authors:** Haley J Fortin, Anna Z Struba, Christopher L Baker

## Abstract

Genetic variation drives phenotypic diversity and disease susceptibility. *Trans*-acting genetic variation coordinates genome-wide chromatin changes, yet the molecular mechanisms underlying this regulation remain unclear. Here, we use the power of mouse genetics to investigate how genetic variation at *trans*-acting loci regulates 3D chromatin interactions.

Using HiChIP to map H3K27ac-associated regulatory elements in C57BL/6J and DBA/2J embryonic stem cells (ESCs), we identified 4,962 strain-differential interactions, 71% of which overlapped chromatin accessibility quantitative trait loci (QTL), establishing chromatin interaction variation is predominantly heritable. These differential interactions showed coordinated changes in chromatin state and gene expression, with stronger interactions associated with increased accessibility and transcription. Notably, loci regulated in *trans* exhibited a unique chromatin signature where weaker interactions were enriched for H3K9me3-marked heterochromatin. Analysis of F1 hybrids revealed dominant repressive effects, consistent with heterochromatin-mediated *trans*-regulation. To causally test this mechanism, we generated reciprocal congenic mouse strains carrying a Chr13 *trans*-QTL region. Integrated multiomic profiling of congenic ESC lines demonstrated that this single locus coordinates changes in H3K9me3, H3K27ac, chromatin accessibility, and 3D contact frequency at hundreds of distal targets. Consistent with our expectations, 73-83% of differential interactions at Chr13 *trans*-QTL targets changed in the predicted direction, demonstrating that heterochromatin-mediated *trans*-regulation coordinately regulates hundreds of regulatory loci.

This work establishes heterochromatin formation as a mechanism by which genetic variation at *trans*-acting loci coordinates changes across chromatin accessibility, histone modifications, and 3D genome organization, providing a framework for understanding how early developmental chromatin states could generate phenotypic variation while preserving essential developmental programs.

## Background

In humans, genetic variation has a profound impact on normal health and disease. The majority of loci associated with variation in complex traits are concentrated in *cis* regulatory elements (CREs), such as promoters, enhancers, and insulators, rather than coding regions [1–3]. Of interest, many of these regulatory elements are active during early development [1,4], suggesting potential developmental origins of adult-onset diseases. The activity of regulatory elements is modulated through DNA and histone modifications, chromatin accessibility, and three-dimensional (3D) genome architecture [5–8]. Given that 99% of the human genome is non-coding, understanding how genetic variation influences CREs is critical for deciphering genome function [9,10], normal development, and, when these processes goes awry, the etiology of developmental disorders and disease.

Sequence variation can alter gene expression locally in *cis*, i.e., on the same chromosome (Chr), or distally, on another chromosome, in *trans*. *Cis*-regulatory variants affect nearby genes through changes in local CREs, whereas *trans*-regulatory effects are mediated by diffusible factors, often transcription factors or chromatin regulators, that can act on many CREs throughout the genome [11]. There is a growing interest in addressing how *trans*-regulation impacts complex traits and disease [12–18]. While effect sizes, tissue-specificity, and multiple testing burden similarly challenge both human and mouse mapping studies [15,16], the reduced power to detect *trans* quantitative trail loci (QTL) in human cohorts is largely driven by allele frequency. Rare variants often exhibit larger effects but require very large sample sizes for detection, whereas common variants, that typically exhibit weaker effects, can be mapped with smaller cohorts [19–21]. In contrast, mapping populations such as the mouse Diversity Outbred (DO) and BXD recombinant inbred panel effectively make rare variants common, enabling more powerful mapping of *trans*-QTL [22–25]. Using the BXD recombinant inbred panel, whose parental strains C57BL/6J (B6) and DBA/2J (D2) differ at over 5,000,000 single nucleotide variants (SNVs) [26], we previously identified both *cis* and *trans* quantitative trait loci (QTL) regulating chromatin accessibility, histone modifications, and gene expression in mouse embryonic stem cells (mESCs) and male germ cells [27,28]. In ESCs, we mapped several *trans-*QTL hotspots on Chrs 4, 5, 7, 12, and 13 that regulate coordinated changes in both chromatin accessibility and gene expression, impacting gene networks important for pluripotency and cell fate determination [28]. Interestingly, genes associated with development and disease have also been independently mapped to closely overlapping intervals within the Chr13 region [27,29–33]. This Chr13 region contains a cluster of KRAB zinc-finger proteins (KZFPs), known transcriptional repressors that act in *trans* to establish heterochromatin. Despite success in identifying *trans*-QTL in both mice and humans [27,28,34–36], their molecular mechanisms remain poorly understood.

Chromosome conformation capture (3C) technologies [37–42] have revealed regulatory roles for 3D genome organization in gene regulation [43–48], from regulatory loops connecting CREs to their target genes [49–52] to topologically associated domains [53–55]. Local genetic variation alters 3D genome structure in *cis*, helping determine how SNVs are associated with disease risk [56] and how structural variants (SVs) impact development of disorders ranging from rheumatoid arthritis and sex reversal to limb malformations [57–59]. Systems genetic approaches have also identified how genetic variation among human lymphoblastoid cell lines influences diverse features of chromatin state and 3D genome architecture [60]. While these studies have established how *cis*-acting genetic variants disrupt 3D chromatin organization, the mechanisms by which *trans*-acting genetic variation influences 3D genome architecture remain largely unexplored. Even when *trans*-QTL are detected, the molecular mechanisms by which single genetic loci coordinate chromatin state changes at hundreds of distal genomic regions are largely unknown. Understanding these mechanisms is essential for interpreting how genetic variation influences complex traits and disease susceptibility, and the contribution of variation to early developmental regulation.

Here, we investigated molecular mechanisms underlying *trans*-regulation of 3D genome organization. Using HiChIP to map H3K27ac-associated chromatin interactions in B6 and D2 mouse embryonic stem cells, we identified that contact anchors of differential interactions substantially overlapped QTL associated with chromatin accessibility (caQTL) and gene expression (eQTL). Integrated epigenomic profiling revealed that heterochromatin formation distinguished *trans* from *cis* regulatory mechanisms. To causally test this mechanism, we performed reciprocal genetic perturbations by exchanging a *trans*-QTL region between otherwise isogenic B6 and D2 genetic backgrounds, enabling independent validation of predicted chromatin changes in both directions. Multiomic profiling of these congenic mESCs validated that this *trans*-acting locus coordinates chromatin state at hundreds of distal regulatory loci with high concordance. Together, these findings establish *trans*-regulation through heterochromatin formation as a key mechanism that coordinates changes in the regulatory landscape.

## Results

### Genetic variation alters chromatin interactions and coordinates gene expression

To investigate how genetic variation impacts 3D genome architecture at CREs, we performed HiChIP [39] on mESC lines derived from B6 and D2 embryos. HiChIP was performed using the histone modification H3K27ac to enrich for 3D interactions at active enhancers. In total, this approach identified a combined set of 90,098 high-confidence interactions across both genetic backgrounds, defined as those spanning greater than 5 kb, a false discovery rate (FDR) < 0.01, and supported by at least four paired-end tag (PET) counts in at least two samples. Independently derived replicate lines from the same strain showed high concordance, while between-strain comparisons revealed reduced concordance, indicating that genetic background, rather than line-specific variation, drives chromatin organization differences (Fig. 1a and Additional file 1: Fig. S1a). These interactions predominantly connected enhancers to promoters (e-p, 84%), or enhancers to other enhancers (e-e, 14%). To understand the functional impact of 3D contacts, we integrated RNA-seq data previously collected from the same ESCs [28]. We observed that gene expression at connected promoters increases with increasing number of chromatin interactions (All pairwise comparisons Tukey p < 2e-16; Fig. 1b), consistent with previous findings [61,62]. Furthermore, genes annotated for “stem cell population maintenance” (MGI GO:0019827) are enriched among the upper quartile of connected genes (Fisher’s Exact Test p < 0.032), consistent with the observation that highly connected genes represent cell type–specific function [63,64]. Among the high-confidence interactions, 4,962 (5.5%) were significantly different between B6 and D2 ESCs (Benjamini-Hochberg corrected p < 0.2; Fig. 1c). To validate HiChIP results, we used 3C-qPCR [65] on three differential interactions (DIs) and confirmed the directional trends of strain-specific interaction patterns (Additional file 1: Fig. S2-4).

**Fig. 1.**
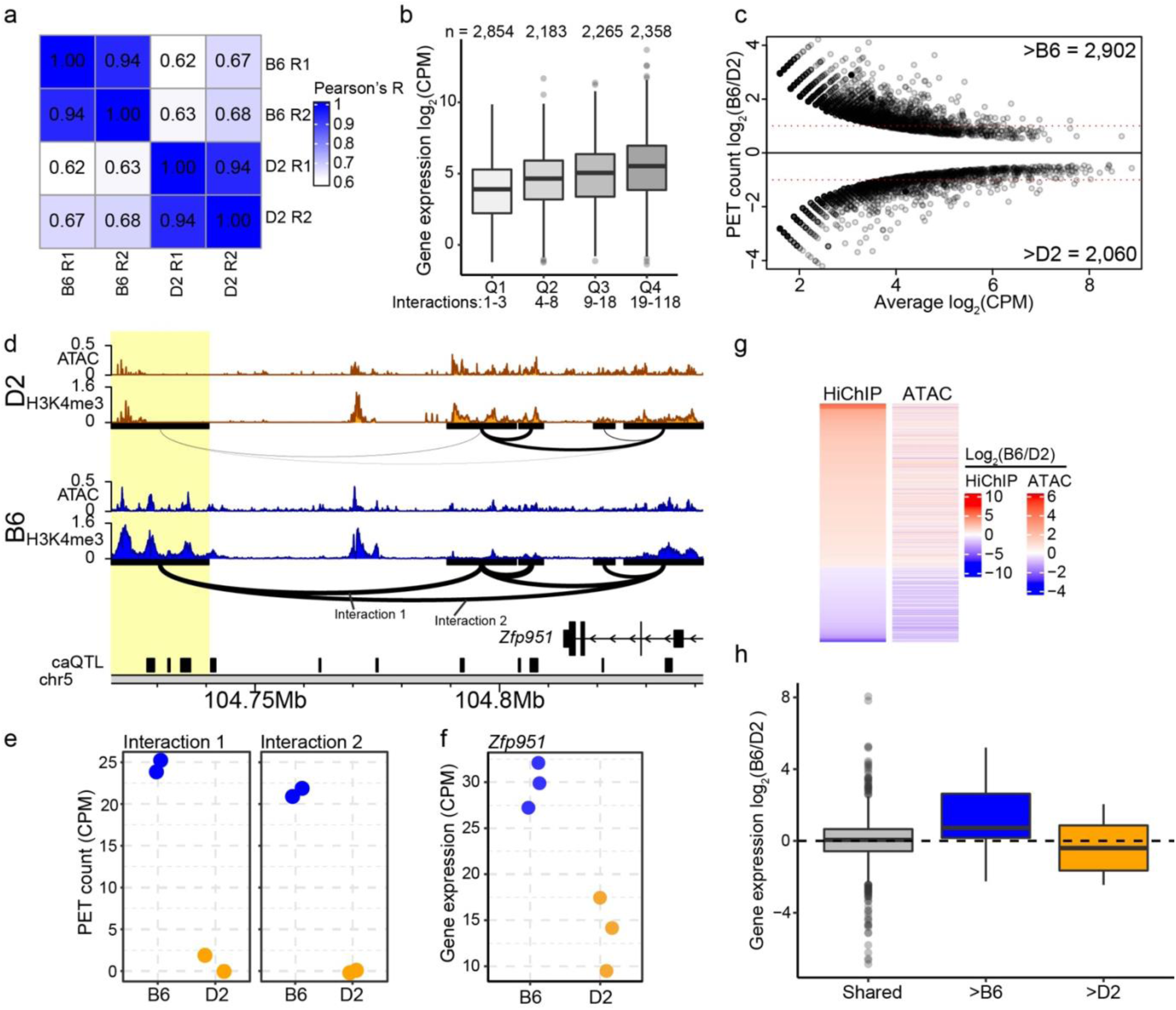
Genetic variation coordinates chromatin interactions, accessibility, and gene expression. **a** Pearson correlation of H3K27ac HiChIP from biological replicates (R1 and R2) of mouse embryonic stem cells (ESCs) derived from B6 and D2 strains. **b** Box and whisker plots indicating mean gene expression values from RNA-seq for B6 and D2 mESCs (for all figure boxes: first quartile, median, and third quartile, whiskers: minimum and maximum, outliers are indicated as dots). Genes were stratified into quartiles (Q1-4) based on the number of contacts per promoter (interactions below graph, gene number above). One-way ANOVA: F_(3, 9656)_ = 232, p < 2e-16; Tukey HSD post-hoc test: all pairwise comparisons between quartiles were significant. **c** MA-plot for differential interactions B6 and D2 ESCs (FDR < 0.2). **d** Genome browser profiles of H3K4me3 ChIP-seq and ATAC-seq at the *Zfp951* locus for B6 (blue) and D2 (orange). DIs are drawn below H3K4me3 tracks. The thickness of the arc connecting two anchors represent contact frequency. Black boxes represent chromatin accessibility regions mapped as quantitative trait loci (caQTL) [28]. Yellow highlight – contact anchor overlapping multiple QTL targets. **e**. HiChIP PET counts for B6 and D2 from interactions 1 (p = 7.44e-9) and 2 (p = 4.48e-10) at *Zfp951* locus. **f** Expression of *Zfp951* from B6 and D2 ESCs (p = 2.81e-5). **g** Heatmap indicating fold-change (B6/D2) in HiChIP PET counts for DIs (left) and corresponding fold-change for ATAC-seq signal at peaks that overlap contact anchors (right). **h** Box and whisker plots of fold-change (B6/D2) in RNA-seq signal at genes which overlap contact anchors for interactions without significant differences (shared, n = 943), and either greater in B6 (n = 17) or D2 (n = 6).

Differential chromatin interactions between B6 and D2 ESCs correspond with coordinated changes in chromatin accessibility and gene expression. The *Zfp951* locus provides an example of coordinated regulation. Specifically, B6 shows higher PET counts at both e-e (Interaction 1) and e-p (Interaction 2) interactions compared to D2 (Fig. 1d, e). Increased 3D contact in B6 is consistent with higher epigenetic marks associated with increased CRE activity (e.g., chromatin accessibility and H3K4me3 enrichment) in the loop anchor region ( Fig. 1D yellow highlight) and higher *Zfp951* expression in B6 (Fig. 1f). This pattern suggests a shared regulatory mechanism coordinating changes in CRE activity, 3D contacts, and gene expression. Genome-wide, DIs showed coordinated strain-specific changes in both chromatin accessibility and interaction strength (Fig. 1g). The strain-specific bias seen for *Zfp951* expression generally holds for all DIs (Fig. 1h); chromatin interactions that are stronger in either B6 or D2 coincide with higher expression of anchor-connected genes. In contrast, genes connected to non-DIs, such as the housekeeping gene *Gapdh*, showed minimal expression bias (log_2_(B6/D2) = 0.05416) (Fig. 1h, Additional file 1: Fig. S1b, c). Together, these data demonstrate that genetic variation coordinately modulates CRE activity, 3D interaction strength, and expression of interacting genes, but leaves open the question of the mechanism of regulation.

### Genetic variation regulates chromatin interactions through both *cis*- and *trans*-acting mechanisms

To test whether coordinated changes in chromatin state and gene expression reflect shared genetic control, we determined if DI anchors are enriched for genetically-regulated CREs by examining overlap with previously identified caQTL targets [28]. For instance, both gene expression of *Zfp951* and accessibility of an upstream CRE (Chr5:104,727,903 bp) were mapped to the same Chr5 locus (Fig. 2a, b, c), with B6 haplotypes conferring higher activity. In total, five interaction anchors contacting *Zfp951* overlap eight *cis*-caQTL all showing higher B6 accessibility [28], demonstrating that individual anchors can harbor multiple caQTL. This pattern extends genome-wide; 71.7% of DI anchors are enriched for caQTL targets at one (42.6%) or both (29.1%) anchors (QTL at one and both anchors p < 0.001, Fig. 2d; Additional file 1: Fig. S1d, e), indicating that contact frequency at anchors for most DIs are under genetic regulation. Further, these data suggest that genetic regulation coordinates both chromatin accessibility and 3D genome structure.

**Fig. 2.**
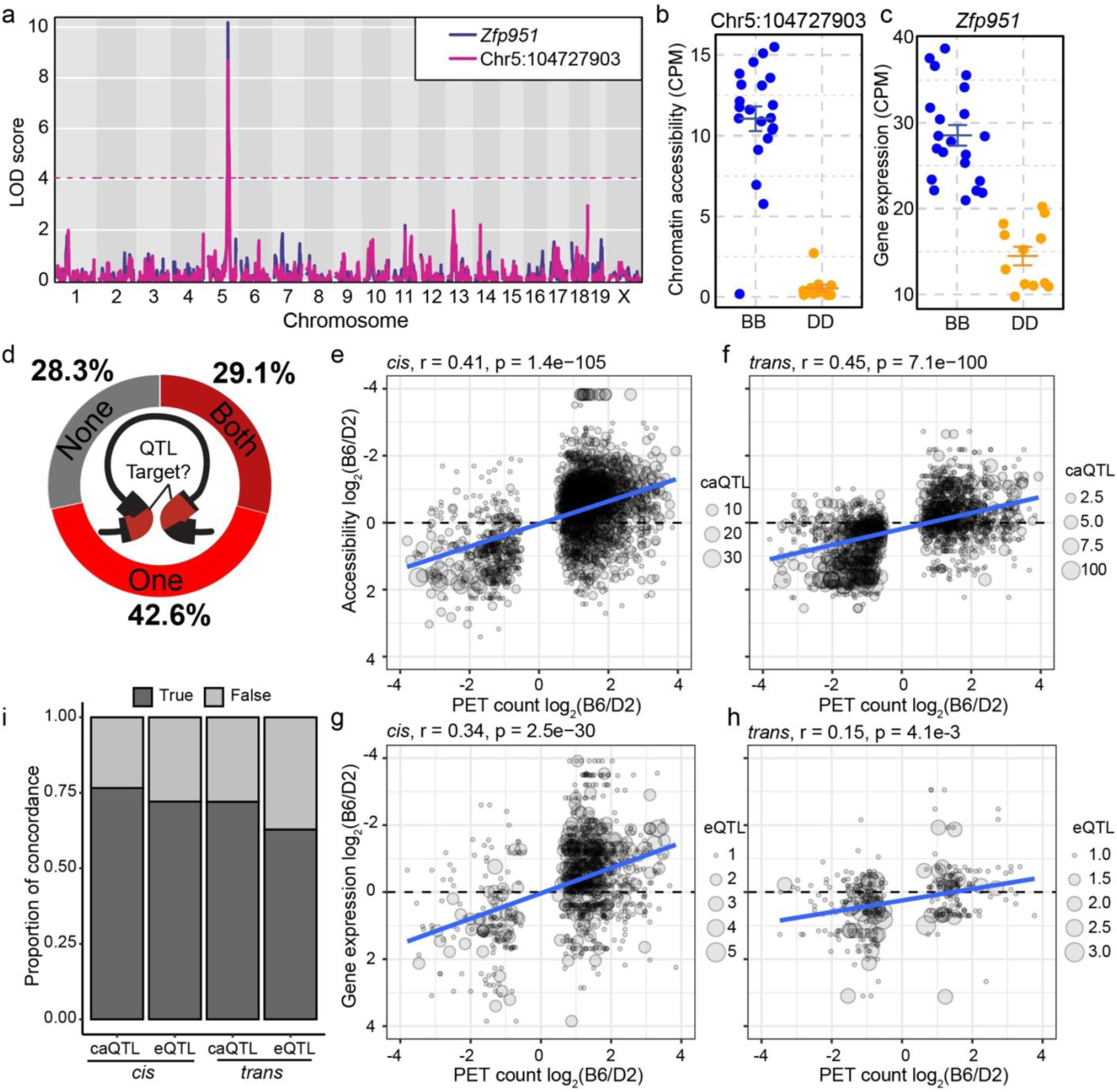
Differential interactions overlap QTL targets and show allelic concordance across regulatory layers. **a** LOD score plot for *Zfp951* and a regulatory element QTL (Fig 1d, leftmost caQTL target in highlighted anchor). Both chromatin accessibility and expression of *Zfp951* are regulated locally on Chr 5. **b** Chromatin accessibility at *Zfp951* regulatory element (Fig 2a pink line) in 33 BXD ESC lines based on genotype at the QTL locus (line represents mean ± standard error). **c** Gene expression of *Zfp951* in 33 BXD ESC lines. **d** Donut plot indicating proportions of DIs for which contact anchors overlap caQTL targets at one, both, or no anchors. Correlation between PET count and chromatin accessibility (**e**, **f**) or gene expression (**g**, **h**), separated into *cis*- (left) and *trans*-associations (right). **i** Proportion of allele concordance between interaction strength and chromatin accessibility (caQTL) or gene expression (eQTL). *cis* caQTL: χ^2^ = 529.5, df = 2, p < 2.2e-16; *trans* caQTL: χ^2^ = 440.2, df = 2 p < 2.2e-16; *cis* eQTL: χ^2^ = 148.2, df = 1 p < 2.2e-16; *trans* eQTL χ^2^ = 20.3, df = 1 = 6.7e-6.

Given QTL enrichment at DI anchors, we tested whether parental interaction strength in B6 and D2 correlates with QTL effect sizes measured across the BXD panel [28]. Interaction strength showed moderate but highly significant positive correlation with chromatin accessibility at *cis* (Pearson r = 0.41, p = 1.4e-105) and *trans* (r = 0.45, p = 7.1e-100) caQTL targets (Fig. 2e, f). We further explored the link between 3D contacts and gene regulation (Fig. 2g, h) and identified a similar correlation at *cis* (r = 0.34, p = 2.5e-30) and *trans* (r = 0.15, p = 4.1e-3) eQTL targets. Beyond correlation magnitude, directional concordance provides stronger evidence for coordinated regulation; loci where the B6 allele shows higher interaction strength might be expected to exhibit B6-biased molecular phenotypes. Indeed, at the categorical level, we found significant allele concordance between interaction strength and chromatin accessibility or gene expression at both *cis-* and *trans*-QTL targets (Fig. 2i, Pearson’s Chi-squared test all p < 6.7e-6). The high allelic concordance between 3D architecture, chromatin accessibility, and gene expression at both *cis*- and *trans*-QTL suggests shared regulatory mechanisms coordinate these molecular layers. Moreover, the presence of *trans*-acting effects indicates that diffusible chromatin regulators mediate long-range genetic control of chromatin organization.

### Heterochromatin formation regulates *trans*-acting chromatin interactions

While both *cis* and *trans*-QTL showed coordinated effects on chromatin accessibility and 3D interactions, the molecular mechanism distinguishing these regulatory modes remained unclear. We hypothesized that distinct mechanisms underlie *trans* and *cis* regulation. To test this, we examined enrichment of DNA-binding factors at interaction anchors that overlap *trans*-regulated loci (i.e., *trans*-targets). Supporting our previous observations [28], we identified TRIM28 (KAP1) as the most highly enriched factor at *trans*-targets (Fig. 3a). Given TRIM28’s role in establishing heterochromatin through H3K9me3 deposition [66,67], we hypothesized that *trans*-targets would show enrichment for repressive chromatin marks. Specifically, at loci with DIs, we predicted H3K9me3 enrichment in the strain with weaker contacts and H3K27ac enrichment in the strain with stronger contacts. To test this, we performed H3K9me3 and H3K27ac ChIP-seq in independently derived mESC lines from three B6 and three D2 mice (Additional file 1: Fig. S5). At a putative CRE downstream from *Sox13* (Fig. 3b, c, d), D2 shows weaker chromatin interactions than B6 and increased H3K9me3 compared to B6 at the anchor overlapping a *trans-*target (Fig. 3b, yellow highlight). Conversely, the same *trans*-target showed increased H3K27ac in B6 compared to D2. To broadly capture this trend, we quantified the level of both H3K27ac and H3K9me3 at both *cis* and *trans*-targets and correlated strain-specific changes in histone modifications with changes in chromatin accessibility (Fig. 3e, f, g, h). When considering H3K27ac, we identified a strong positive correlation at *cis*- (Pearson r = 0.83, p < 2.2e-16) and *trans-* (r = 0.91, p < 2.2e-16) caQTL targets, indicating that both signify active chromatin. The key distinction in chromatin state is with H3K9me3. While *cis-*targets showed little correlation between H3K9me3 and accessibility (Fig. 3g), *trans-*targets showed a strong negative correlation (r = −0.66, p = 5.8e-143; Fig. 3h), indicating that heterochromatin formation specifically characterizes *trans*-regulation. Critically, *trans*-targets were significantly more likely than *cis*-targets to show discordant chromatin states – defined as opposing changes in accessibility and H3K9me3 enrichment (p < 2.2e-16, odds ratio = 20.49, logistic regression; Fig. 3i), establishing heterochromatin formation as a defining feature of *trans*-regulation. Together, these data support a molecular mechanism in which genetic variation in *trans*-acting factors drives TRIM28-mediated heterochromatin formation via H3K9me3 deposition, restricting chromatin accessibility and weakening 3D contacts at distal loci.

**Fig. 3.**
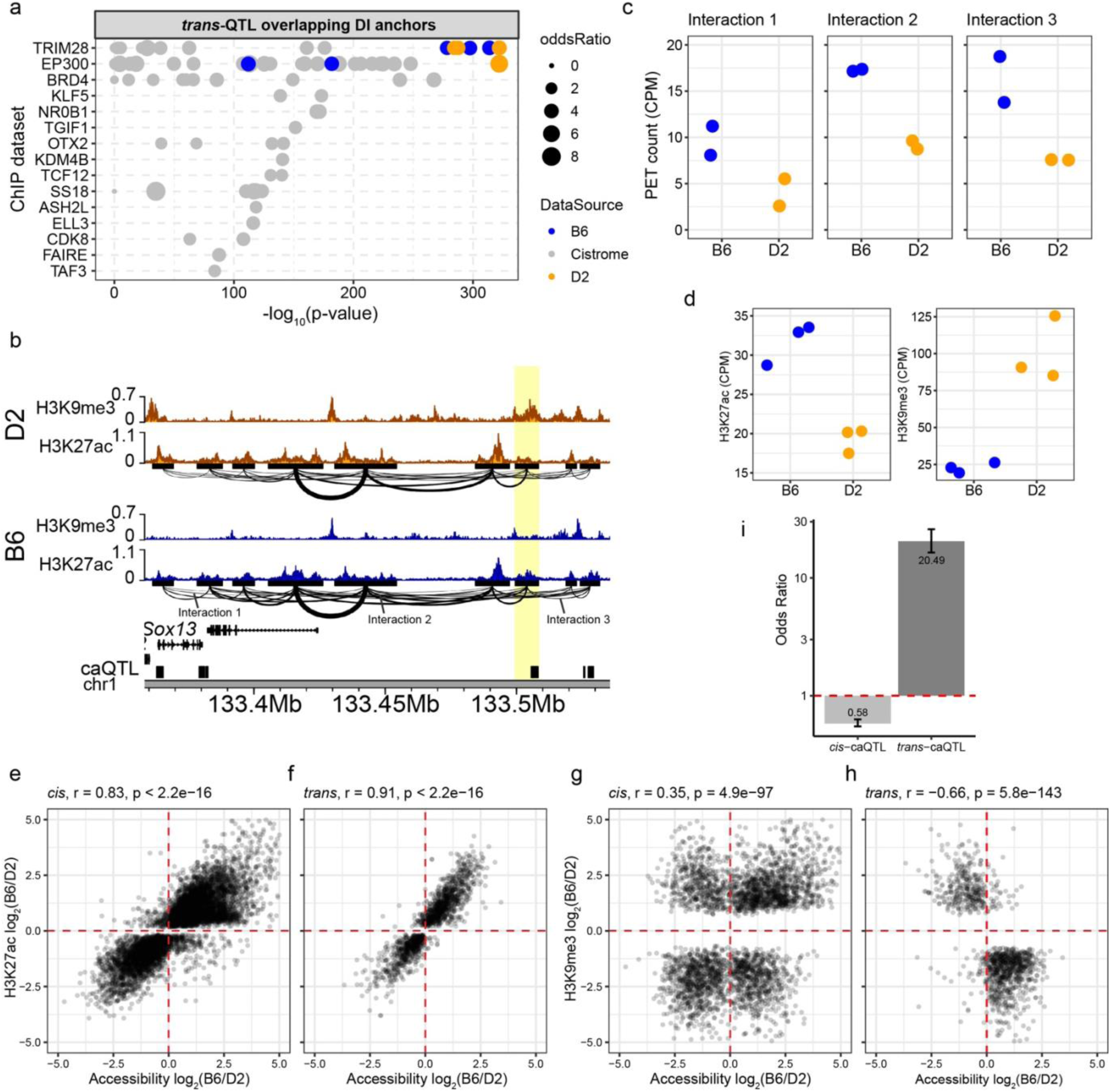
Heterochromatin formation distinguishes *trans* from *cis* regulatory mechanisms. **a** Enrichment of the top 15 DNA-binding proteins at *trans*-QTL targets overlapping DI anchors. Each dot represents a ChIP-seq experiment either from publicly available database (grey - cistrome), or our previously published B6 (blue) and D2 (orange) datasets. Binding sites were evaluated for overlap between *trans*-targets and DI contact anchors (Fisher’s exact test, top 15 factors with q-value cutoff < 0.01). **b** Genome profiles for H3K27ac and H3K9me3 ChIP enrichment from D2 (orange) and B6 (blue) at *Sox13* locus. HiChIP interactions indicated below H3K27ac tracks. Highlight indicates a contact anchor that overlaps a *trans*-target (black box). **c** HiChIP PET counts for three significant DIs indicated in panel *b* (Interaction 1 p = 9.53e-3, Interaction 2 p = 1.05e-2, Interaction 3 p = 3.04e-3). **d** Difference in H3K27ac (left) and H3K9me3 (right) at caQTL target highlighted in panel *b* between B6 and D2 (H3K27ac p = 2.24e-3, H3K9me3 p = 3.58e-3). Correlation between fold change (B6/D2) in H3K27ac enrichment and chromatin accessibility at *cis-* and *trans*-QTL targets (**e, f**). Correlation between fold change in H3K9me3 enrichment and chromatin accessibility at *cis-* and *trans*-QTL targets (**g, h**). **i** Odds ratio showing that *trans*-targets are 20.49 times more likely to show discordant chromatin states resulting in high H3K9me3 and low ATAC.

### Chr13 *trans*-QTL coordinates chromatin accessibility and gene expression in an allele-specific manner

To determine the effect of *trans*-regulation on coordinated chromatin regulation and gene expression, we focused on a compound *trans*-QTL on Chr 13 that regulated the expression of 59 genes (eGenes) and 2,033 putative CREs (*trans*-targets) with LOD > 5 [28]. Spatial coordination of *trans*-targets and eGenes was assessed for proximity enrichment using a modified Region Associated Differentially expressed gene approach [68], testing whether chromatin accessibility peaks cluster near transcriptionally regulated genes. Chr13 *trans*-targets showed higher enrichment near Chr13 eGenes compared to all other *trans*-QTL eGenes (Fig. 4a), indicating distal *trans*-regulation drives coordinated changes in chromatin accessibility and gene expression at co-localized target loci. Critically, this spatial coordination was directionally concordant with QTL effects; chromatin peaks with higher accessibility associated with the B6 allele at the QTL are specifically enriched near eGenes with higher expression when the genotype at the QTL is B6 (Fig. 4b, left), and the reciprocal pattern held for D2 (Fig. 4b, right). These analyses demonstrate that independently measured chromatin accessibility and gene expression are spatially coordinated in an allele-specific manner, supporting *cis*-propagation of *trans*-QTL through localized chromatin remodeling.

**Fig. 4.**
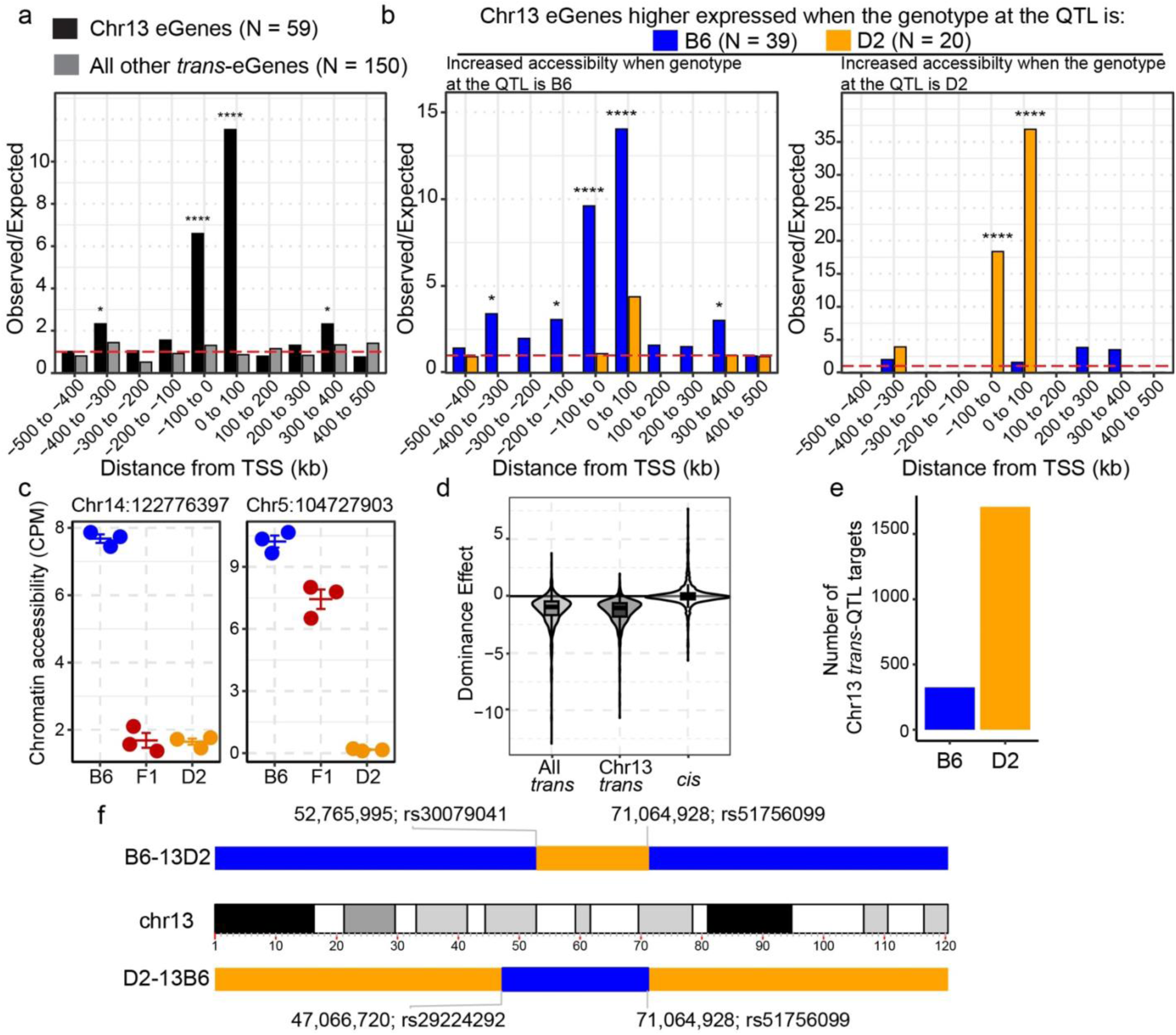
Chr13 *trans*-QTL coordinates spatial regulation of chromatin accessibility and gene expression through dominant repression. **a** Observed-over-expected ratio for the spatial proximity of Chr13 *trans*-targets to transcription start sites (TSS) of either Chr13 eGenes (black) or all other *trans*-eGenes (grey). Red dashed line is null expectation (binomial test, * p < 0.05, ** p < 0.01, *** p < 0.001, **** p < 0.0001). **b** Similar to *a* except stratifying genes that either higher expressed when Chr13 QTL haplotype is B6 (blue) or D2 (orange). Spatial proximity was tested for peaks with higher chromatin accessibility when Chr13 is B6 (left) or D2 (right). **c** Scatterplot of chromatin accessibility based on B6, D2, or F1-hybrid genotype of ESCs at two caQTL. Left – representative Chr13 *trans*-target indicating dominant repressive effect on chromatin accessibility. Right - *cis*-QTL target representative of additive effect (line – mean, bars – standard error). **d** Distribution of dominance effect values in all *trans*-targets (N = 836), Chr13 *trans*-targets (N = 575), and all *cis*-caQTL (N = 7049). Negative values indicate repressive effect; values near zero indicate additive effect. **e** Classification of Chr13 *trans*-targets by repressive allele. To establish directional predictions targets were stratified by which QTL genotype reduces chromatin accessibility, consistent with dominant repression. **f** Schematic of chromosome 13 showing the base-pair position and identifying SNP for congenic regions in B6-13D2 (D2 region on B6 background) and D2-13B6 (B6 region on D2 background) strains.

To determine the nature of the QTL effect on chromatin accessibility, we performed ATAC-seq on three independently derived mESCs from (B6xD2)F1 hybrids. Focusing on high-confidence caQTL (LOD > 8), *trans*-caQTL showed dominant repression, where F1 hybrid mESCs resembled the parent with lower accessibility rather than intermediate (additive) values. This is exemplified by the Chr13 *trans*-target at Chr14:122776397, where the F1 showed the same (dominant) repressed accessibility as the D2 parent (Fig. 4c, left). In contrast, a *cis*-QTL target at Chr5:104727903 showed additive inheritance with F1 chromatin accessibility intermediate between the parent lines (Fig. 4c, right). This distinction between *trans* and *cis* effects extended genome-wide; *trans*-caQTL showed dominant repression (median r = −0.95), while *cis*-caQTL showed near-additive inheritance (median r = −0.04; Wilcoxon p < 2.2e-16, effect size r = 0.61; Fig. 4d). Chr13 *trans*-caQTL targets specifically showed strong dominant repression (median = -1.04), with the D2 allele at the Chr13 QTL associated with reduced accessibility at most targets (84%, N = 1,709) (Fig. 4e).

The dominant repressive inheritance pattern supports a model in which the Chr13 QTL encodes one or more diffusible repressors that establish heterochromatin at *trans*-targets. To causally test whether this locus coordinates heterochromatin formation and 3D architecture, we generated reciprocal congenic mouse strains that exchange the Chr13 *trans*-QTL. The first strain, B6.D2-(rs30079041-rs51756099)/Bakr (hereafter referred to as B6-13D2), was bred to contain the D2 Chr13 congenic region (Chr13:52,765,995-71,064,928) on an otherwise B6 background. Conversely, D2.B6-(rs29224292-rs51756099)/Bakr (hereafter referred to as D2-13B6) was bred to carry the B6 Chr13 congenic region (Chr13:47,066,720-71,064,928) on an otherwise D2 background (Fig. 4f). Together, these congenic lines provide ideal resources to investigate the impact of the Chr13 *trans*-QTL on heterochromatin formation.

### Congenic strains confirm Chr13 *trans-*QTL drives heterochromatin formation at *trans*-targets

To test whether the Chr13 QTL drives *trans*-regulation of heterochromatin, we tested predictions based on QTL effects of BXDs on reciprocal mESCs from congenic strains. If Chr13 is causal, swapping the QTL region should alter chromatin state according to the genotype introduced. Specifically, B6-13D2 should resemble D2 at *trans*-targets, while D2-13B6 should resemble B6. We tested these predictions using H3K9me3 and H3K27ac ChIP-seq, and ATAC-seq in B6, D2, B6-13D2, and D2-13B6 mESCs (Additional file 1: Fig. S6).

If repression is mediated by H3K9me3-marked heterochromatin, we predicted *trans*-targets should show increased H3K9me3 in B6-13D2 (with D2 Chr13) compared to B6. Strikingly, 77% of Chr13 *trans*-caQTL targets showed the predicted H3K9me3 increase in B6-13D2 (Fig. 5a, g, left; exact binomial test p < 2.2e-16). Reciprocally, 89% showed decreased H3K9me3 in D2-13B6 (with B6 Chr13) compared to D2 (Fig. 5b, left, g; p < 2.2e-16). Of the remaining targets (324 loci predicted to be repressed when the allele at the QTL is B6), 69% showed the inverse pattern where H3K9me3 increased in D2-13B6 and decreased in B6-13D2 (Fig. 5b, c, right, g). Consistent with H3K9me3-mediated repression, active chromatin marks showed the reciprocal pattern as predicted. Targets gaining H3K9me3 in B6-13D2 showed reduced chromatin accessibility (77% of predicted decreases validated; Fig. 5c, g) and reduced H3K27ac (94% validated; Fig. 5e, g). Similarly, targets losing H3K9me3 in D2-13B6 gained accessibility and H3K27ac (Fig. 5d, f, g). Further, *trans*-targets that changed in the predicted direction showed significantly larger effect sizes than those that did not (Fig. 5h), indicating non-validated targets likely reflect measurement noise rather than biological exceptions. These reciprocal epigenomic changes, validated through two independent genetic perturbations across three chromatin features, directly demonstrate that Chr13 causally regulates hundreds of distal *trans*-targets through heterochromatin formation.

**Fig. 5.**
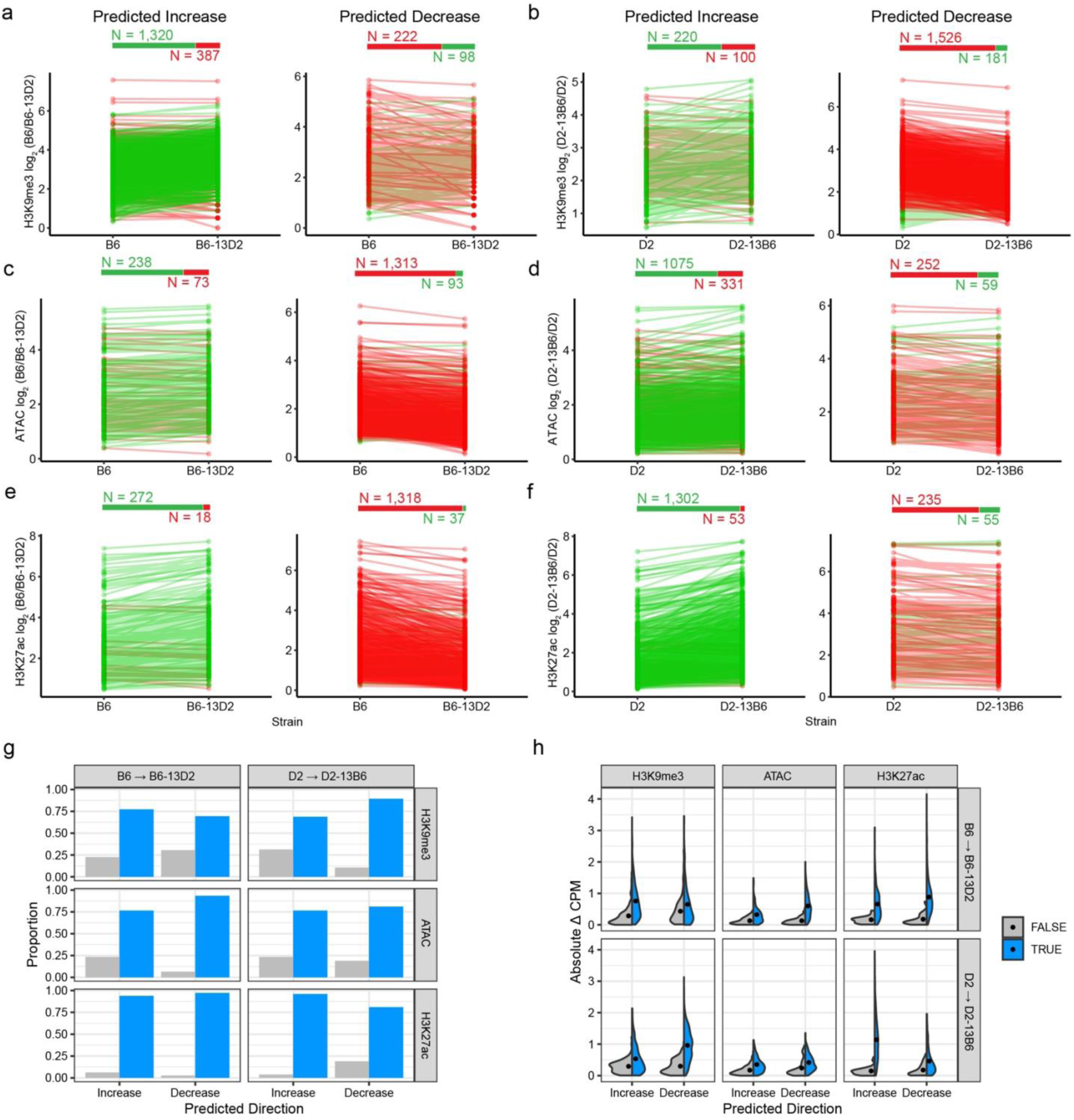
Chr13 *trans*-QTL regulates heterochromatin formation at hundreds of distal loci. **a-f** Histone modification or chromatin accessibility at Chr13 *trans*-targets in parental (B6 and D2) and reciprocal congenic (B6-13D2 and D2-13B6) strains for H3K9me3 ChIP-seq (**a, b**), ATAC-seq (**c, d**), and H3K27ac ChIP-seq (**e, f**). Each line represents a single *trans*-target, connecting measurements between the parental strain (left) and its corresponding congenic (right). B6→B6-13D2 panels (**a, c, e**) test the effect of introducing the D2 Chr13 region into the B6 background, while D2→D2-13B6 panels (**b, d, f**) test the reciprocal introduction of the B6 Chr13 region into the D2 background. Within each panel, loci are separated by predicted direction of change based on QTL effect direction from BXD mapping: "Predicted Increase" (left) indicates targets expected to gain signal in the congenic strain, while "Predicted Decrease" (right) indicates targets expected to lose signal. Green lines indicate increasing values, while red lines indicate decreasing values. Numbers above each plot show total *trans*-targets tested that increase (green) or decrease (red). Statistical significance assessed by exact binomial test. H3K9me3 enrichment: **a** increase p < 2.2e-16, decrease p = 3.3e-12; **b** increase p = 1.7e-11, decrease p < 2.2e-16 ATAC-seq: **c** increase p < 2.2e-16, decrease p < 2.2e-16; **d** increase p < 2.2e-16, decrease p < 2.2e-16. H3K27ac: **e** increase p < 2.2e-16, decrease p < 2.2e-16; **f** increase p < 2.2e-16, decrease p < 2.2e-16. **g** Observed proportion of Chr13 *trans*-targets for which chromatin state changes in the direction predicted (i.e., True) based on QTL effect and dominant repression. **h** Violin plot showing distribution of absolute changes in counts per million (CPM) at Chr13 *trans*-QTL targets. *Trans*-targets that change as predicted (blue) or do not change as predicted (grey). Dots indicated median value.

### Heterochromatin formation alters 3D contacts at *trans*-QTL target sites

To validate that Chr13-driven heterochromatin formation alters 3D interactions at *trans*-targets, we repeated H3K27ac HiChIP in B6 and D2, while including B6-13D2 and D2-13B6 mESCs (two technical replicates per strain; Additional file 1: Fig. S7). We hypothesized that introducing the D2 Chr13 region in B6-13D2 would reduce chromatin interaction at *trans*-targets (compared to B6), by increasing heterochromatin. Further, introducing the B6 Chr13 region into D2-13B6 would strengthen interactions (compared to D2), by reducing heterochromatin. Consistent with our hypothesis, at the *Litaf* locus we observed increased interactions in D2-13B6 compared to D2 (Fig. 6a, b, Interactions 1 and 2 p < 0.027). Reciprocally, at the same locus we observe weaker interactions in B6-13D2 compared to B6 (Fig 6a, b; Interactions 1 and 2, p < 1.51e-3). This is particularly noteworthy because *Litaf* is a Chr13 eQTL target, highlighting a potential functional link between chromatin architecture and gene regulation.

**Fig. 6.**
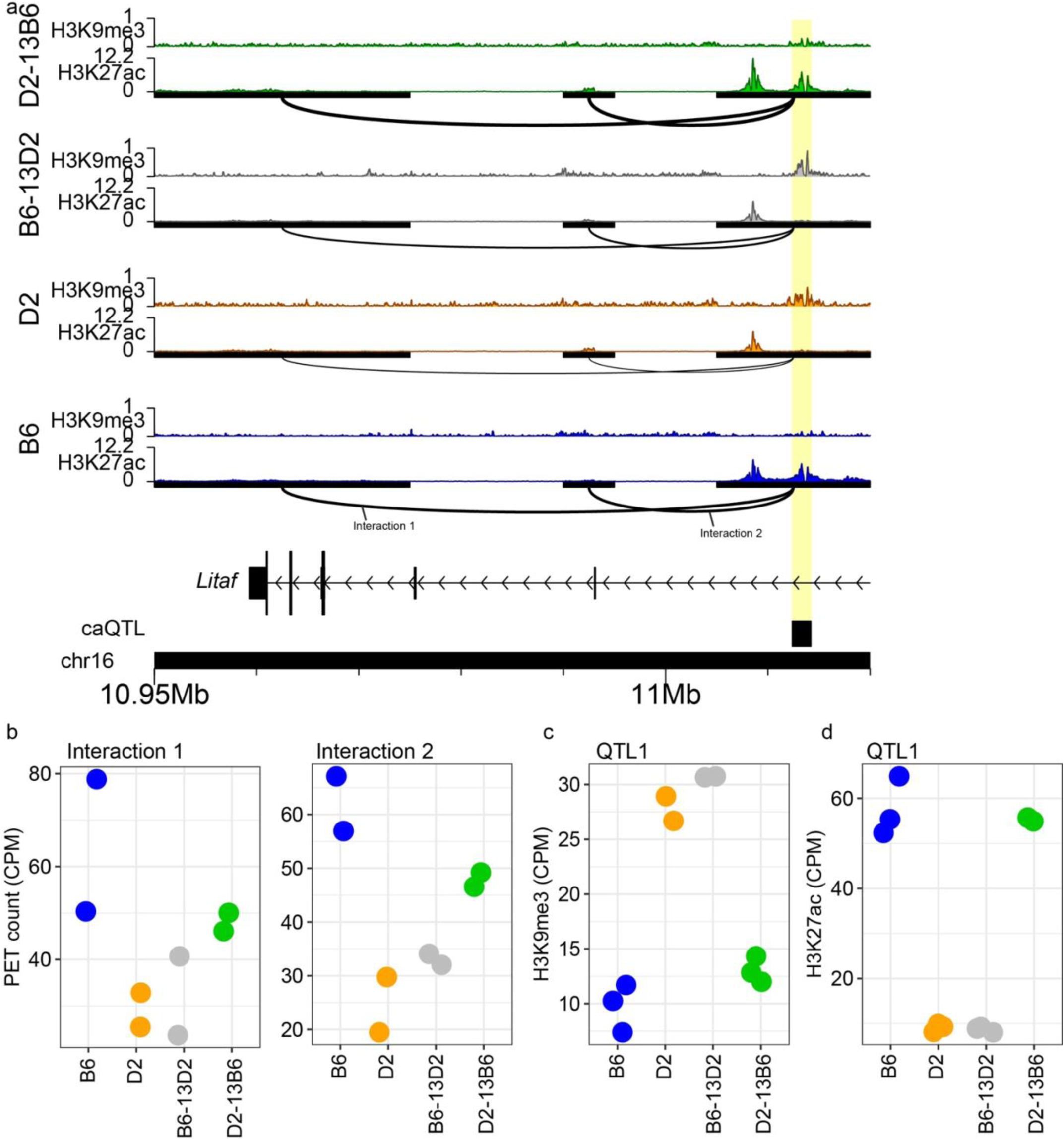
Chr13 coordinates heterochromatin and 3D architecture at *Litaf* locus. **a** Genome profiles for H3K27ac and H3K9me3 ChIP enrichment from B6 (blue), D2 (orange), B6-13D2 (grey) and D2-13B6 (green) at the *Litaf* locus. HiChIP interactions indicated below H3K27ac tracks with the thickness of the arc connecting two anchors representing contact frequency. Highlight indicates a *trans*-target (black box) overlapping a contact anchor and variable histone modifications. **b** HiChIP PET counts for all four strains for two interactions indicated in panel a (Interaction 1: BD p = 4.67e-4, BB13D p = 3.14e-4, DD13B p = 2.68e-2; Interaction 2: BD p = 9.94e-5, BB13D p = 1.51e-3, DD13B p = 3.15e-3). **c** H3K9me3 enrichment at Chr13 *trans-*target highlighted panel a (BD p = 8.25e-12, BB13D p = 2.98e-15, DD13B p = 4.81e-8). **d** H3K27ac enrichment at Chr13 *trans* caQTL target highlighted panel a (BD p = 6.62e-108, BB13D p = 4.54e-119, DD13B p = 9.24e-103). BD is the comparison between B6 and D2; BB13D is the comparison between B6 and B6-13D2; DD13B is the comparison between D2 and D2-13B6.

To test if heterochromatin formation underlies these changes in chromatin interactions, we examined H3K9me3 enrichment at the interaction anchors. Critically, H3K9me3 showed the reciprocal pattern to 3D contacts. At *Litaf*, D2-13B6 exhibited significantly lower H3K9me3 than D2 at the Chr13 *trans*-target; and B6-13D2 showed increased H3K9me3 compared to B6 (Fig. 6c; DD13B p = 4.81e-8, BB13D p = 2.98e-15). These examples demonstrate that the Chr13 QTL drives heterochromatin formation at interaction anchors, coincident with altered 3D interaction strength. Further, in agreement with a heterochromatin-mediated mechanism, H3K27ac showed the opposite patterns, with D2-13B6 exhibiting higher levels at the same anchor compared to D2 at *Litaf* (Fig. 6d; p < 9.24-103), while B6-13D2 showed lower H3K27ac than B6 (Fig 6d; BB13D p = 4.54e-119). Consistent DI patterns and opposing H3K9me3/H3K27ac profiles are observed at additional *trans*-regulated loci, including *Zfp516* and *Fam241a* (Additional file 1: Fig. S8-9). Together these data confirm that Chr13 regulates heterochromatin formation and 3D organization at these *trans*-targets.

To assess whether these examples reflect genome-wide patterns, we tested whether genetic differences identified between parental strains (B6 and D2) predict changes in congenic strains. Remarkably, 73-83% of differential interactions changed in the predicted direction (Fig. 7a, b, c; Exact binomial test: all p < 1.5e-8) Furthermore, interactions that change contact frequency as predicted generally showed larger effect sizes than those that did not (Fig. 7d), indicating non-validated targets may reflect measurement noise. These validation rates, achieved across two independent genetic perturbations and testing reciprocal predictions, demonstrate that Chr13 causally regulates chromatin state at hundreds of distal loci. The consistent validation across H3K9me3, H3K27ac, chromatin accessibility, and 3D contact frequency establishes *trans*-regulation of heterochromatin as a mechanism for coordinated regulation of the epigenome (Fig. 7e).

**Fig. 7.**
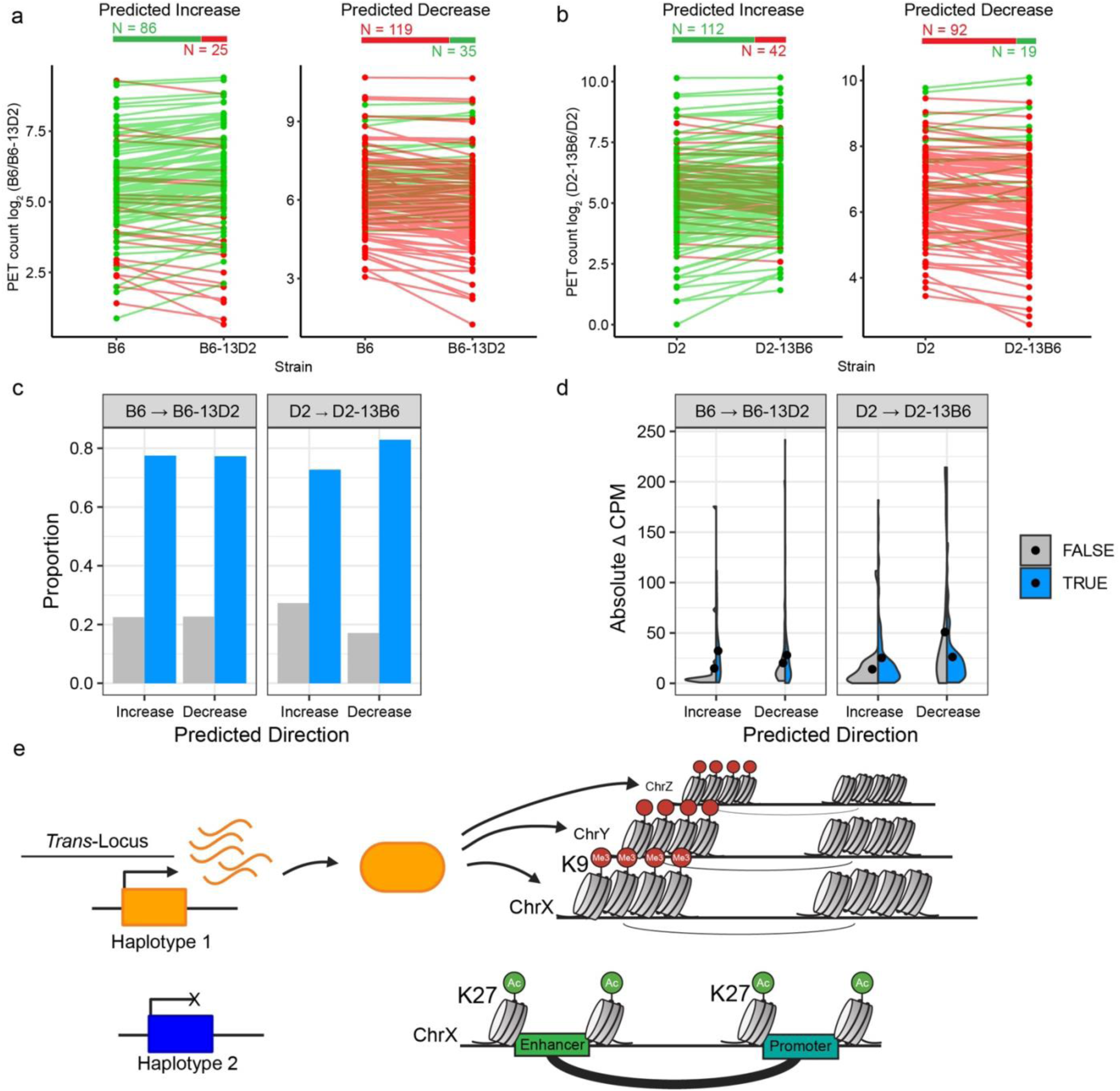
Heterochromatin coordinates genome-wide changes in 3D architecture. **a**, **b** Validation of predicted changes in 3D contact frequency at Chr13 *trans*-targets using H3K27ac HiChIP. Experimental design and visualization as in Fig. 5a-f, with each line representing a single interaction overlapping a Chr13 *trans*-target. B6→B6-13D2 (**a**) and D2→D2-13B6 (**b**). Loci were split by predicted direction of change based on BXD QTL mapping. Green lines indicate increasing PET counts; redlines indicate decreasing PET counts. Numbers above show total interactions tested that increase (green) or decrease (red). Exact binomial test: **a** increase p = 5.03e-9, decrease p = 6.59e-12; **b** increase p = 1.53e-8, decrease p = 1.12e-12. **c** Observed proportion of interactions that changed as predicted (blue). **d** Distribution of absolute changes in PET counts at Chr13 *trans*-targets, showing validated predictions (blue) and non-validated targets (grey). Dots indicate median values. **e** Model illustrating *trans*-regulation through heterochromatin formation. In this example, the orange haplotype *encodes* diffusible repressor(s) that establish H3K9me3-marked heterochromatin at distal *trans*-targets (ChrX, Y, Z), reducing chromatin accessibility and weakening 3D contacts. Whereas the blue haplotype lacks active repressor(s), permitting open chromatin and stronger enhancer-promoter interactions and gene expression at the same distal locus.

## Discussion

Here, we establish heterochromatin formation as the molecular mechanism distinguishing *trans* from *cis* regulatory effects on 3D genome organization in mESCs. *Trans*-regulated regions of the genome show a 20-fold enrichment for discordant H3K9me3 and chromatin accessibility, providing a unique epigenetic signature compared to *cis*-QTL. This mechanism is supported genetically by the dominant repressive pattern observed in F1 hybrids. Critically, reciprocal genetic perturbations demonstrated that a single locus causally coordinates chromatin state at hundreds of distal regulatory elements with high concordance across molecular assays. Validation rates ranged from 73-83% for 3D contact frequency to 89-94% for H3K9me3 and H3K27ac, demonstrating that the coordinated mechanism is robust across the epigenome. Together, these findings explain how *trans*-acting genetic variation achieves coordinated regulation across molecular layers through targeted heterochromatin deposition.

Consistent with previous observations [60,62,69,70], we find that 3D genome interactions are positively correlated with gene expression and active histone modifications, with contact frequency at individual loci associated with increased transcription. Prior work in human induced pluripotent stem cells (iPSCs) demonstrated that genetic variation affecting contact propensity results in coordinated changes in H3K27ac [71,72]; however, these studies did not distinguish *cis* from *trans* regulation. Our findings extend beyond correlations by establishing causal relationships through reciprocal genetic perturbations and identifying heterochromatin formation as a defining feature of *trans*-regulation. Notably, the 20-25% of cases where increased contacts did not correspond to increased expression suggest additional regulatory mechanisms beyond direct connection between contact frequency and expression and warrant future investigation.

The functional genomic, genetic, and positional evidence continue to point to variation in genes that encode KRAB Zinc-Finger Proteins (KZFPs) as probable molecular basis for *trans*-regulation. Firstly, KZFPs provide DNA-binding specificity to recruit TRIM28, and chromatin modifying enzymes, to form heterochromatin [66,73–77]. Consistent with this, TRIM28 is enriched at *trans-*QTL targets and a cluster of KZFPs is found in the Chr13 QTL [78]. Second, the unique association of *trans*-QTL with H3K9me3 and their dominant repressive effect in F1 hybrids indicates a diffusible silencing factor [27,28]. Third, KZFPs are highly variable within species, indicating a source of rapid genetic variation [78,79]. The Chr13 region represents a compound QTL containing structurally variant KZFP genes, each encoding distinct DNA-binding specificities; this genetic architecture explains how a single locus coordinates changes at hundreds of *trans*-targets. Fourth, KZFPs have been demonstrated to impact transcriptional repression through heterochromatin spreading from their DNA binding sites [80], supporting the regional *cis*-acting effects we see on enhancer-promoter contacts. Lastly, the epigenetic engineering approach of CRISPR interference (CRISPRi) [81] leverages a nuclease-dead Cas9 fused to a KRAB domain to induce heterochromatin and silence target genomic loci, directly demonstrating *trans*-regulation of heterochromatin. Building on this, recent work used CRISPRi to show that targeted H3K9me3 deposition led to a reduction in 3D contacts between enhancers and promoters [69], demonstrating direct effect of heterochromatin on 3D contacts in an inducible system.

We propose that *trans*-regulation of heterochromatin formation represents an evolutionarily advantageous mechanism for generating regulatory variation without disrupting core transcription factor (TF) networks. Classical models of *trans*-regulation invoke changes in TF expression or sequence, which would coordinately affect all targets genome-wide. However, developmental TFs and their core gene regulatory networks are under strong purifying selection due to their essential roles in lineage specification ; and large variation in these factors likely results in severe phenotypic consequences. In contrast, KZFP DNA-binding domains evolve rapidly through structural variation [82,83], and specifically at Chromosome 13 in mice [78], potentially enabling population-level variation in which genomic regions are differentially targeted for heterochromatin formation based on the repertoire of zinc-finger domains present in an individual. Recent cross-species comparisons between human and macaque cells demonstrate that *trans*-regulatory divergence is specifically enriched at transposable elements bearing KZFP footprints [84], providing evolutionary support that heterochromatin variation drives regulatory divergence between populations while preserving core developmental networks. Further, heterochromatin acts as a barrier to pioneer transcription factor binding during cellular reprogramming [85,86], and depletion of a specific KZFP reduced this barrier and enhanced reprograming efficiency [87]. The coordinated regulation we observe here reveals a powerful and underappreciated layer of genome control where a single locus affects heterochromatin deposition and therefore chromatin accessibility and 3D contacts at hundreds of regulatory elements. By allowing population-level variation in heterochromatin targeting rather than disrupting essential TF function, this mechanism may provide standing regulatory variation necessary for evolutionary adaptation while maintaining developmental robustness.

The developmental implications of heterochromatin-mediated *trans*-regulation established in early pluripotency may extend to adult phenotypes. Heterochromatin can act as a barrier to lineage specification [85]; and variation in early heterochromatin formation may be propagated, creating a persistent epigenetic state that influences lineage commitment and cellular function. Supporting functional consequences, the Chr13 region investigated here has been associated with multiple developmental phenotypes in mice including craniofacial development, limb malformation, lupus susceptibility, meiotic recombination frequency, and imprint stability [27,29–33]. The mechanism we establish in mouse ESCs appears conserved in human biology. Inter-individual variation in heterochromatin landscapes across diverse human iPSC lines correlates with differentiation propensity for early neural fates versus other lineages [88], consistent with the model that heterochromatin variation creates tunable barriers affecting lineage-specific transcription factor access. This suggests population-level variation in heterochromatin may contribute to phenotypic diversity in humans through the same mechanism. Further, KZFPs show dynamic expression beyond pluripotency, specifically upregulated in adult immune cells and neurons [83]. Consistent with functional relevance, genomic regions containing KZFPs in humans contain signatures of recent selection and are associated with adult disease [89]. Together, these observations suggest that early variation in heterochromatin establishment may act to constrain developmental outcomes and contribute to phenotypic variation in complex traits.

Our study provides strong genetic and epigenetic evidence that heterochromatin formation mediates *trans*-regulation of 3D genome organization and chromatin state, but several limitations remain. First, all experiments were performed in mESCs, and whether similar mechanisms operate in differentiated tissues, other species, or in vivo is unknown. Second, the Chr13 congenic interval spans a large region and contains multiple KZFPs, making it unclear whether *trans*-regulation reflects the combined effects of this region or a single causal gene. Fine-mapping, and targeted perturbations, such as overexpression or CRISPR-based manipulation, will be needed to identify the causal gene(s). Finally, while reciprocal congenic experiments validate chromatin changes, functional consequences for pluripotency, lineage specification, and organismal phenotypes remain to be tested; and extension to human systems will be essential to establish relevance to complex traits and disease.

## Conclusions

We establish heterochromatin formation as a defining chromatin signature distinguishing *trans* from *cis* regulatory mechanisms in mouse embryonic stem cells. Through reciprocal genetic perturbations, we demonstrate that genetic variation at a single *trans*-acting locus regulates chromatin accessibility, histone modifications, and 3D interactions at hundreds of distal regulatory elements. This work provides a framework for understanding how *trans*-regulation operates mechanistically through targeted heterochromatin formation to reshape regulatory landscapes and suggests that genetic variation in early developmental chromatin states may contribute to phenotypic diversity in complex traits and disease susceptibility.

## Methods

### Mice

All mice were obtained from The Jackson Laboratory (Bar Harbor, ME) including C57BL/6J (JR# 000664) and DBA/2J (JR# 000671) parental inbred strains. B6 and D2 parents were crossed to the F3 generation prior to marker-based genotyping and subsequent backcrossing for a minimum of ten generations for form the B6.D2-(rs30079041-rs51756099)/Bakr (JR# 37111) and D2.B6-(rs29224292-rs51756099)/Bakr (JR# 37112) congenic lines. Congenic genotypes were confirmed using MiniMUGA array [90] (Neogen) for which the background analysis report is available for a representative mouse (B6-13D2: Additional file 2; D2-13B6: Additional file 3). All animal experiments were approved by the Animal Care and Use Committee of The Jackson Laboratory (Summary #04008). Congenic lines are available upon request.

### Derivation and maintenance of mouse embryonic stem cells

Derivation of mouse embryonic stem cells for B6-13D2 and D2-13B6 strains were performed, after backcrossing for a minimum of 10 generations, using protocols outlined in Czechanski et al. [91]. For all cultures, mouse ESCs were seeded with irradiated mouse embryonic fibroblasts (MEFs) from B6 animals and cultured in DMEM high glucose base medium supplemented with 15% fetal bovine serum (FBS, Lonza, cat. no. 14-501F lot no. 0000217266), 1X Pen/Strep, 2 mM GlutaMAX, 1 mM sodium pyruvate, 0.1 mM MEM-NEAA, 0.1 mM 2-mercaptoethanol, 103 IU LIF, 1 µM PD0325901, and 3 µM CHIR99021. Mouse ESCs were expanded for 2–3 days to reach ∼70% confluency for molecular assays. For functional genomic assays, mouse ESCs were enzymatically disassociated into single cell suspensions using trypsin and MEFs were depleted by incubation on gelatine coated plates for 20 minutes to allow to settle. All cell lines tested negative for mycoplasma and are available upon request. Genotyping was performed to determine zygosity of congenic region and sex of cell lines. Primers for KASP genotyping used SNVs within the congenic region to distinguish between B6 and D2 backgrounds. All primers used for genotyping are listed in Additional file 4, Table S1.

### ATAC-seq sample preparation and sequencing

ATAC-seq experiments were performed, using two or three independent cultures from the same cell line per strain, following the FAST-ATAC method as described [92], using 100,000 cells. DNA quality and quantity was determined using High Sensitivity D5000 ScreenTape (Agilent Technologies). Libraries were sequenced to a read depth of about 40 million reads per sample on an Illumina NovaSeq X Plus with a 150 bp read length.

### Chromatin Immunoprecipitation (ChIP) sample preparation and sequencing

ChIP-seq experiments performed with B6 and D2 used three independently derived cell lines from each strain. Validation ChIP-seq experiments performed with B6, D2, B6-13D2, and D2-13B6 used two or three independent cultures from the same cell line for each strain. For each ESC ChIP-seq library, 3 million cells were harvested and fixed as previously described [27]. Cell lysis, chromatin fragmentation, and immuno-precipitation were performed as described [93], with minor modifications. For H3K27ac ChIP, sodium butyrate was added to hypotonic lysis buffer, MNase buffer, RIPA buffer (post-overnight antibody conjugation), RIPA wash buffer, and TE to inhibit histone deacetylase activity. Immunoprecipitation was performed using antibodies against H3K27ac (12 μL, abcam, ab4729, lot# GR312658-1; Cell Signaling Technology, 8173, lot# 9) and H3K9me3 (6 μL, Active Motif, 39161, lot# 09919003). ChIP-seq libraries were constructed using the KAPA HyperPrep Kit (Roche Sequencing and Life Science). Quality and concentration of libraries were assessed using the High Sensitivity D5000 ScreenTape (Agilent Technologies) and KAPA Library Quantification Kit (Roche Sequencing and Life Sciences). Libraries were sequenced to a read depth of about 40 million reads per sample (75 bp) on an Illumina NextSeq or NovaSeq X Plus.

### HiChIP sample preparation and sequencing

#### Restriction enzyme based HiChIP

HiChIP experiments were performed as described [39], using two samples from independently derived cell lines per strain. We used 10 million cells, DpnII (NEB, R05443), and the following 3C modifications [94] to improve digestion, incorporation, ligation, and removal of dangling ends. Specifically, 1) biotin incorporation was performed at 23°C for 4 hours with interval shaking (10 sec at 900 rpm, 5 min off). 2) Ligation was performed at 16°C for 4 hours with interval shaking (1 min at 500 rpm, 4 min off). 3) Samples were sheared using a Diagenode Biorupter with the following parameters: tube size = 1.5 mL, time on = 30 sec, time off = 30 sec, cycle # = 8. 4) A total of 3 µL of H3K27ac antibody (Abcam #ab4729) was added to the sample and incubated at 4°C overnight with rotation. 5) After ChIP washes, samples were resuspended in 20 µL removal of biotin-dATP master mix: 5 µL 10X NEB Buffer 2.1, 0.125 µL of dATP and dGTP at 10 mM each, 5 µL T4 DNA polymerase (NEB, M0203), and 39.8 µL distilled water. Samples were incubated at 20°C for 4 hours with interval shaking (1 min at 400 rpm, 4 min off). Lastly, size selection of libraries was performed using a two-sided selection with AMPureXP beads. Briefly, 22.5 µL of beads were added to the sample to capture fragments less than 700 bp. Supernatant was transferred to a new tube and 13.5 µL of beads were added to the supernatant sample to capture fragments greater than 300 bp. Libraries were eluted in 50 µL 10 mM Tris pH 8.0. Libraries were quantified using qPCR against Illumina primers and then paired end sequenced with read length of 75 bp to a sequencing depth of 200 million reads per sample over two Illumina NextSeq runs.

#### Dovetail HiChIP MNase Kit

The Dovetail HiChIP MNase Kit was used for HiChIP for validation experiments repeating B6, D2, and including B6-13D2 and D2-13B6. HiChIP was performed using two samples from independent cultures from the same cell line per strain. The User Guide [95] (version 2.0) was used to process samples of 10 million cells. H3K27ac (Cell Signaling Technology, 8173, lot #9) was used for immunoprecipitation. Libraries were paired end sequenced with a read length of 150 bp to a sequencing depth of 300 million reads per sample (over two flow cell lanes) on an Illumina NovaSeq X Plus.

### ATAC-seq and ChIP-seq data processing

Illumina adaptors were trimmed from ChIP reads using Trimmomatic [96] (version 0.39) and then aligned to mouse reference genome (mm10), modified to incorporate known D2 variants reported in Mouse Genomes Project Database, REL-1505 [26,97] using hisat2 [98] (version 2.2.1). Duplicate reads were removed using Picard Tools [99] (version 2.0.1), and peaks were called for each sample using MACS [100] (version 1.4.2). While independent peak calling was performed on H3K27ac and H3K9me3 to determine quality and reproducibility, to compare ATAC- and ChIP-seq signal at QTL targets the comprehensive set of open chromatin locations (i.e., peakome) previously developed [28] was used to generate read counts across all samples using bedtools [101] (version 2.9.2) similar to ATAC-seq. Read matrices were TMM-normalized and log_2_-transformed downstream differential analysis of histone modification using edgeR [102] (version 3.34.1). For visualization on genome browser tracks with karyoploteR [103] (version 1.20.3) we used bwa [104] (version 0.7.5; filter mapping quality < 30, unmapped, secondary, QC fail, duplicates, and supplementary reads) alignment to generate bigwig files.

### HiChIP data processing and analysis

#### Restriction enzyme-based HiChIP

HiChIP reads were processed using HiC-Pro [105] (v2.11.4) with the following parameters:

N_CPU = 4, SORT_RAM = 768M

REFERENCE_GENOME = mm10

LIGATION_SITE = GATCGATC

MIN_FRAG_SIZE = 30

MAX_FRAG_SIZE =100000

MIN_INSERT_SIZE = 30

MAX_INSERT_SIZE = 600.

Long-range interactions were called with hichipper [106] (v.0.7.8b0) using H3K27ac ChIP-seq peaks generated in this paper. PET counts were imported into RStudio (version 4.1) using diffloop [107] (v1.22.0). Before filtering data, read matrices were TMM-normalized. Anchors of interactions were not merged and interactions less than 5,000 bp were filtered out. P-values were adjusted using an FDR cutoff of 0.01. Interactions were filtered for at least four read counts in at least two samples before identifying differential interactions using DESeq2 [108] within the diffloop [107] package. Significant differential interactions were cutoff at FDR < 0.2.

Differential interactions (DIs) showed higher average PET counts than non-differential interactions (Wilcoxon rank sum test p < 2.2e-16; Additional file 1: Fig. S10), consistent with increased statistical power to detect differences at well-covered loci.

#### Dovetail HiChIP MNase Kit

HiChIP reads were processed using Dovetail’s HiChIP Loop Calling [109] (readthedocs Revision 8f2e0b16).

Reads were aligned to mm10 using bwa [104] (v0.7.18). pairtools [110] (version 1.1.2) was used to record and sort valid ligation events, remove PCR duplicates, and generate .pairs and bam files. Samtools [111] (version 1.19.2) was used to create and index the final bam file. Libraries were verified to have no-dup read pairs > 50%, about 20% or greater of the total mapped non-dup pairs to be in *cis* and > 1 kb in length, and total reads in 1 kb around center of ChIP peaks > 2%. Filtered pairs were converted to Hi-C Pro valid pairs format. Loops were called using FitHiChIP [112] (version 11.0) with the following parameters in the bias correction/coverage bias configuration file:

Interaction type = peak to peak Bin size = 2500

Lower distance threshold = 10000

Upper distance threshold = 2000000

Bias types = coverages bias regression

Merge filtering was employed (option 1)

q-value = 0.01

Chromosome sizes correspond to mm10.

Peak-to-peak interactions were imported into RStudio (version 4.1). diffloop [107] (version 1.22.0) was used to adjust p-values using an FDR cutoff of 0.01. Interactions were filtered for at least four read counts in at least two samples. PET counts were normalized, and edgeR [102] (version 3.34.1) was used to identify differential interactions (FDR < 0.2).

### Spatial Enrichment Analysis

Region Associated Differentially expressed gene (RAD) framework [68] was adapted to test whether chromatin accessibility changes and gene expression changes regulated by *trans*-QTL spatially co-localize. The genomic background was defined as genes expressed in ESCs consistent with cutoff used for QTL mapping [28], rather than all annotated genes genome-wide. Strand-specific signed distances between each gene’s TSS to nearby chromatin accessibility peaks were calculated, with negative values indicating peak positions upstream of the TSS in the direction of transcription and positive values indicating downstream positions. Distances were binned into 100 kb intervals spanning -500 kb to +500 kb relative to the TSS. Binomial tests were used to determine significance treating each gene-peak connection within 500 kb as an independent trial (binom.test in R, alternative = ’greater’), with the expected probability defined by the genome-wide proportion of focal genes (p = M_focal / N_total), and Benjamini-Hochberg correction was applied for multiple testing within each analysis.

### 3C-qPCR

3C-qPCR experiments were performed as previously described [65] with the following modifications. Briefly, 1) 10 million cells were fixed with fresh formaldehyde at a final concentration of 1% for 10 minutes with rotation before quenching the reaction with 10X glycine for 5 minutes with rotation. Cells were pelleted, washed with PBS, and flash frozen; 2) 800 U restriction enzyme (NEB, #R0101) and 1 mM ATP were used for digestion; 3) 4,000 U ligase (NEB, #M0202) was used for an overnight incubation at 16°C; 4) Qiagen MaXtract HD tubes were used for 3C sample purification. BACs overlapping loci of interest (Additional file 4, Table S2) were used for PCR control templates. Primers used for 3C-qPCR experiments are listed in Additional file 4.

## Declarations

### Ethics approval and consent to participate

All animal procedures were approved by the Animal Care and Use Committee of The Jackson Laboratory under protocol number 04008.

### Availability of data and materials

Original data discussed in this study in NCBI’s Gene Expression Omnibus and are accessible through GEO Series accession number GSE312918 (https://www.ncbi.nlm.nih.gov/geo/query/acc.cgi?acc=GSE312918).

Along with raw sequencing data, processed data tables in the accession include raw read counts for ATAC-seq, H3K27ac ChIP-seq, H3K9me3 ChIP-seq, and HiChIP. Additionally, there are bigwig files for each ATAC, ChIP, and HiChIP sample. H3K4me3 ChIP-seq [27], ATAC-seq, and RNA-seq [28] for B6 and D2 were collected previously and are available through GEO accessions (H3K4me3: GSE113192, https://www.ncbi.nlm.nih.gov/geo/query/acc.cgi?acc=GSE113192; ATAC-seq and RNA-seq: GSE164935, https://www.ncbi.nlm.nih.gov/geo/query/acc.cgi?acc=GSE164935).

## Supporting information

MiniMUGA Genotyping Results for B6-13D2

MiniMUGA Genotyping Results for D2-13B6

## Acknowledgements

We would like to thank all the members of the Baker laboratory for comments and discussion. Additionally, we would like to thank Steve Munger for comments on the manuscript and Natalie Powers and Petko Petkov for their early breeding of the congenic lines. We recognize contributions from The Jackson Laboratory Genome Technologies services. The Jackson Laboratory scientific services are supported in part through the National Cancer Institute’s Cancer Core Grant P30CA034196.

## Authors’ contributions

CLB and HJF conceptualized the study and contributed to methodology. CLB provided resources. HJF and AZS investigated the study. HJF and CLB performed data curation and formal analysis. HJF and CLB performed visualization. HJF and CLB wrote the manuscript. HJF and CLB revised and edited the manuscript. CLB performed supervision. CLB performed funding acquisition.

