## Supplementary Figures S1-S10 for "*Trans*-regulation of heterochromatin underlies genetic variation in 3D genome contacts"

### Additional File 1 (Supplementary Figures S1 to S10)

---

**Fig. S1 HiChIP differential interactions are enriched for caQTL targets.**

**Fig. S2 3C-qPCR validates stronger interactions in B6 at *Zfp951* locus.**

**Fig. S3 3C-qPCR validates stronger interactions in D2 at *Bicc1* locus.**

**Fig. S4. 3C-qPCR validates stronger interactions in B6 at *Manbal* locus.**

**Fig. S5. Quality control of ChIP-seq data reveals strong reproducibility among replicates.**

**Fig. S6. Quality control of ChIP-seq and ATAC-seq reveals strong reproducibility among replicates.**

**Fig. S7. Quality control of HiChIP data reveals strong reproducibility among replicates.**

**Fig. S8. Chr13 coordinates heterochromatin and 3D architecture at *Fam241a* locus**.

**Fig. S9. Chr13 coordinates heterochromatin and 3D architecture at *Zfp516* locus**.

**Fig. S10. Differential interactions show higher PET counts than non-differential interactions.**

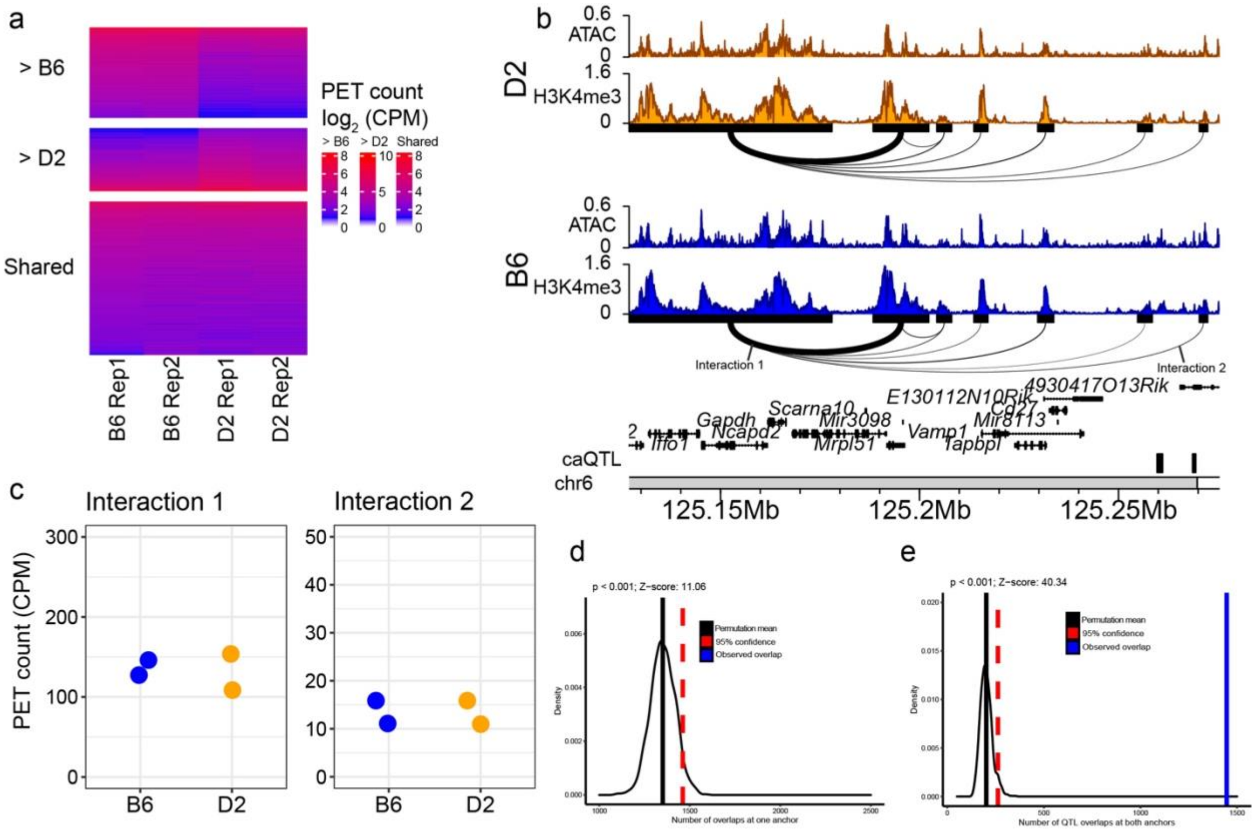


**Fig. S1. HiChIP differential interactions are enriched for caQTL targets.**
**a** Heatmap indicating $\log_2$CPM in HiChIP PET counts for DIs stronger in B6 (> B6), stronger in D2 (> D2), and non-differential interactions (Shared). Heatmap shows interactions from a random sampling of 2,500 DIs (> B6 N = 1,472, > D2 N = 1,028) and 2,500 non-DIs. **b** Genome browser profiles of H3K4me3 ChIP-seq and ATAC-seq at the *Gapdh* locus for B6 and D2. DIs are drawn below H3K4me3 tracks. The thickness of the arc connecting two anchors represent contact frequency. **c** HiChIP PET counts for two non-DIs indicated in panel *b*. **d** Density plot showing enrichment of caQTL targets overlapping one or both (**e**) anchors of DIs.

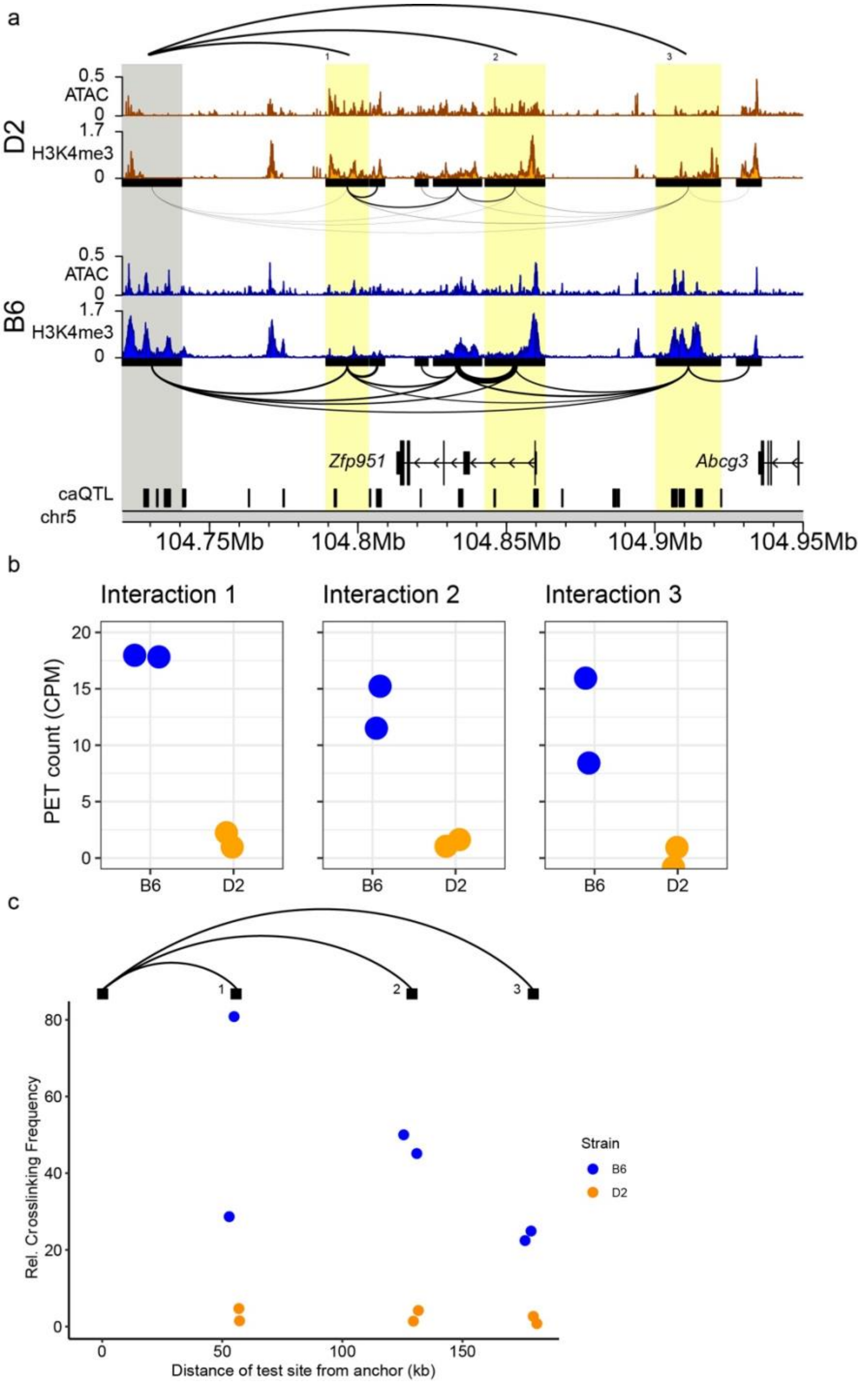


**Fig. S2. 3C-qPCR validates stronger interactions in B6 at *Zfp951* locus.**
**a** Genome profiles for H3K4me3 ChIP-seq and ATAC-seq at the *Zfp951* locus. HiChIP interactions indicated below H3K4me3 tracks. Grey and yellow highlights indicate the anchor and test sites used for 3C-qPCR, respectively. **b** HiChIP PET counts for three significant DIs indicated in panel *a* (Interaction 1 p = 1.65e-12, Interaction 2 p = 2.28e-10, Interaction 3 p = 5.64e-11). **c** Relative crosslinking frequency of interactions at *Zfp951* locus in two biological replicates from B6 and D2 ESCs. Anchor site indicated in panel *a* originates at 0 kb and test sites are located at 55 kb (p = 3.00e-1), 128 kb (p = 4.07e-3), and 177 kb (p = 4.34e-3).

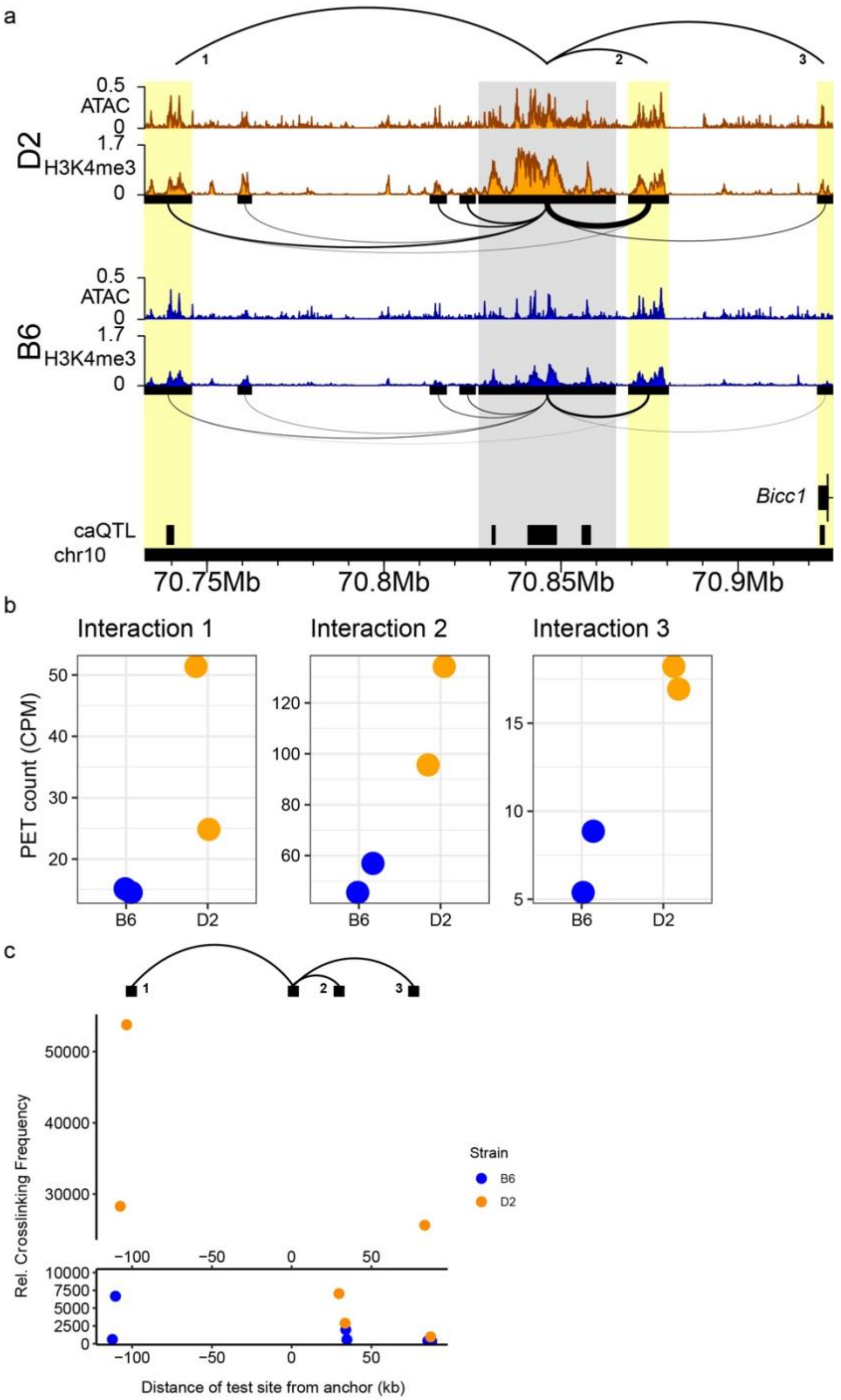


**Fig. S3. 3C-qPCR validates stronger interactions in D2 at *Bicc1* locus.**
**a** Genome profiles for H3K4me3 ChIP-seq and ATAC-seq at the *Bicc1* locus. HiChIP interactions indicated below H3K4me3 tracks. Grey and yellow highlights indicate the anchor and test sites used for 3C-qPCR, respectively. **b** HiChIP PET counts for three significant DIs indicated in panel *a* (Interaction 1 p = 1.81e-4, Interaction 2 p = 2.11e-5, Interaction 3 p = 2.71e-2). **c** Relative crosslinking frequency of interactions at *Bicc1* locus in two biological replicates from B6 and D2 ESCs. Anchor site indicated in panel *a* originates at 0 kb and test sites are located at -108 kb (p = 1.94e-1), 33 kb (p = 3.07e-1), and 83 kb (p = 4.85e-1).

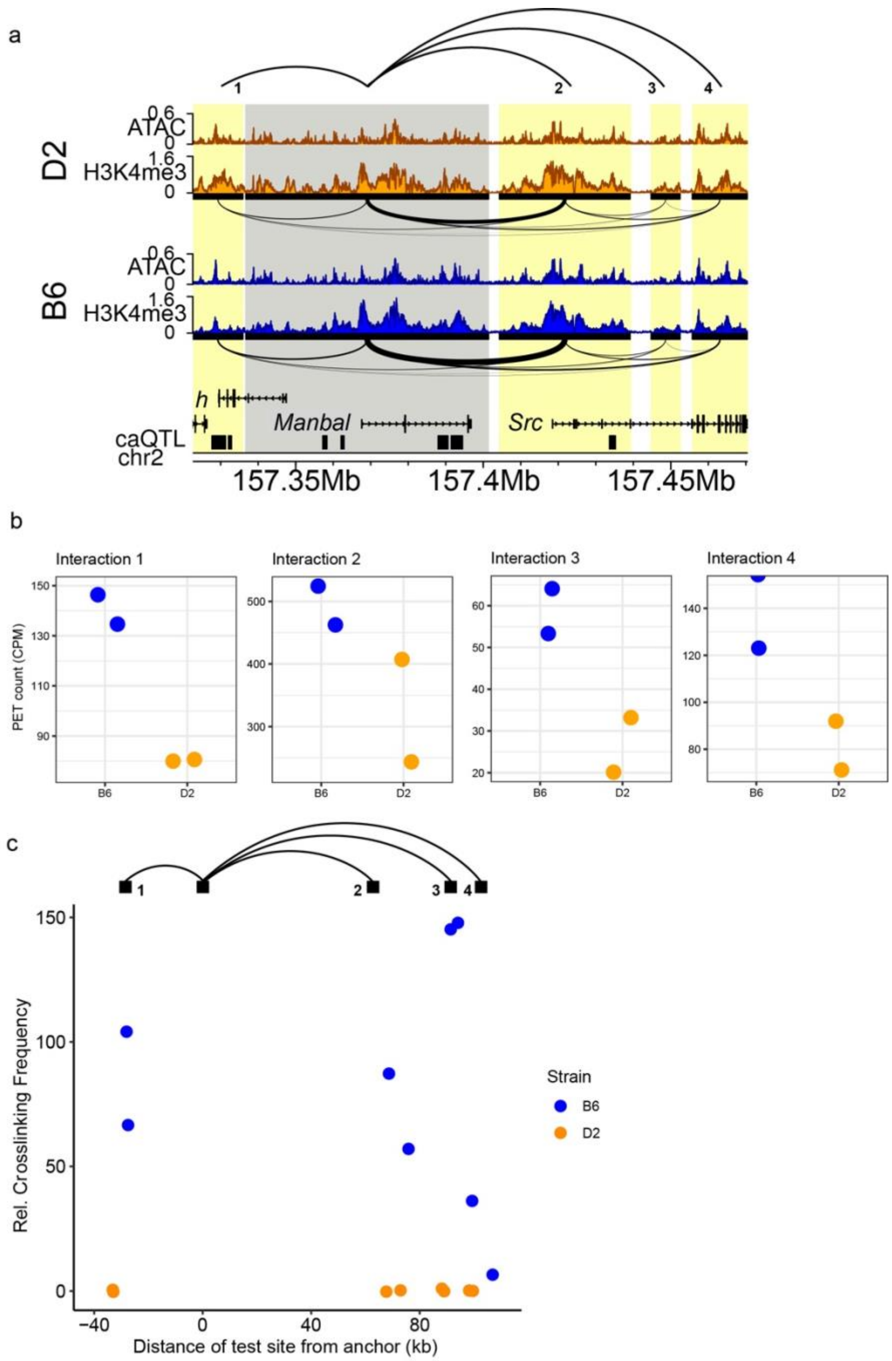


**Fig. S4. 3C-qPCR validates stronger interactions in B6 at *Manbal* locus.**
**a** Genome profiles for H3K4me3 ChIP-seq and ATAC-seq at the *Manbal* locus. HiChIP interactions indicated below H3K4me3 tracks. Grey and yellow highlights indicate the anchor and test sites used for 3C-qPCR, respectively. **b** HiChIP PET counts for four significant DIs indicated in panel *a* (Interaction 1 p = 5.23e-5, Interaction 2 p = 3.39e-4, Interaction 3 p = 9.69e-6, Interaction 4 p = 2.71e-3). **c** Relative crosslinking frequency of interactions at *Manbal* locus in two biological replicates from B6 and D2 ESCs. Anchor site indicated in panel *a* originates at 0 kb and test sites are located at -31 kb (p = 1.36e-1), 72 kb (p = 1.30e-1), 90 kb (p = 3.94e-3) and 103 kb (p = 3.82e-1).

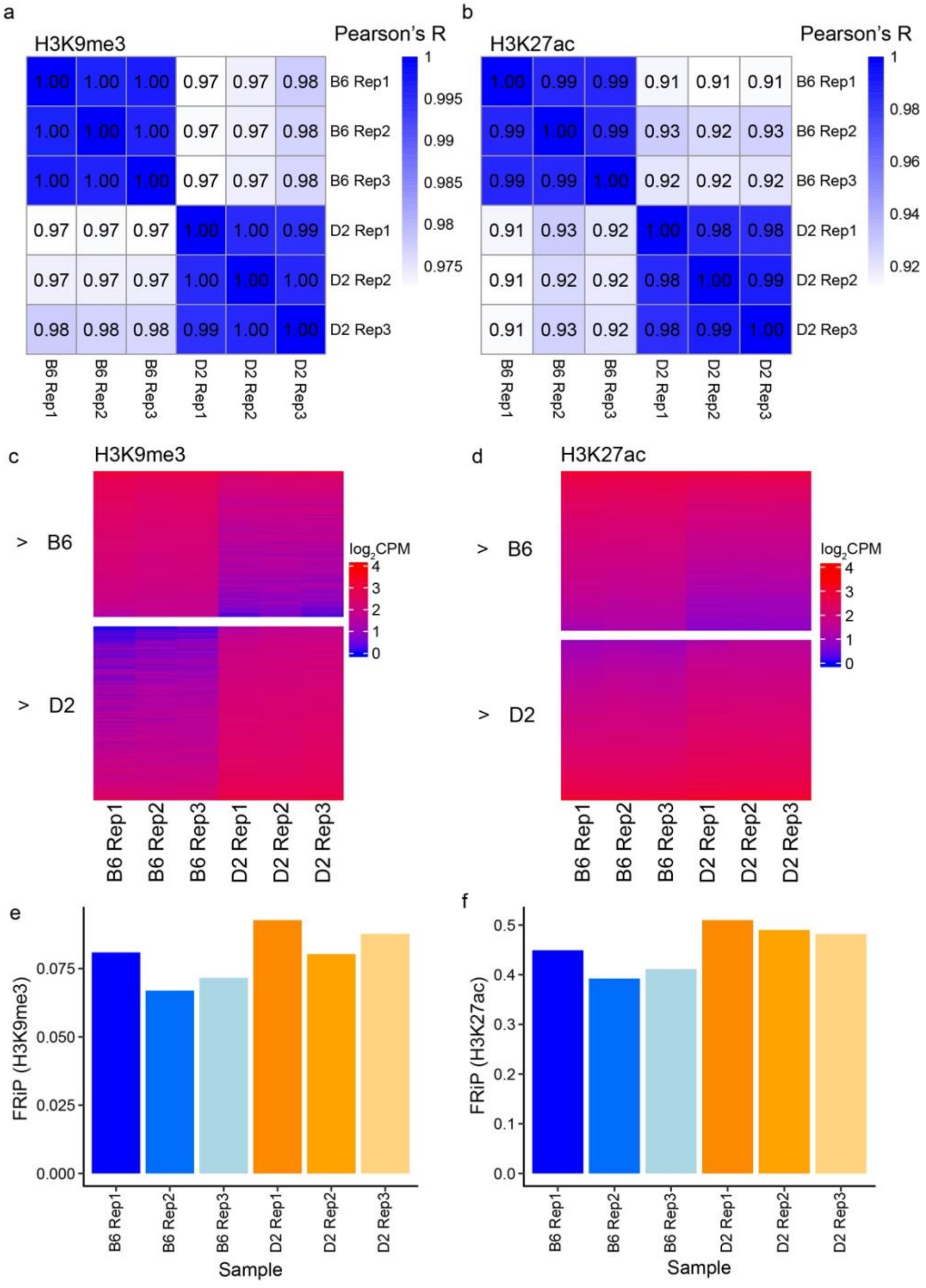


**Fig. S5. Quality control of ChIP-seq data reveals strong reproducibility among replicates.** **a** Pearson correlation of H3K27ac and H3K9me3 (**b**) ChIP from biological replicates of mouse embryonic stem cells (ESCs) derived from B6 and D2 strains. **c** Heatmap indicating $\log_2$CPM in H3K9me3 ChIP read counts for peaks more enriched in B6 (N = 6,391) and more enriched in D2 (N = 7,591). **d** Heatmap indicating $\log_2$CPM in H3K27ac ChIP read counts from a random sampling of 2,500 H3K27ac peaks that are more enriched in B6 or in D2. **e** Fraction of reads in peaks for H3K9me3 and H3K27ac (**f**) ChIP-seq. For H3K9me3, peaks are those identified as ATAC-seq peaks in BXD population.

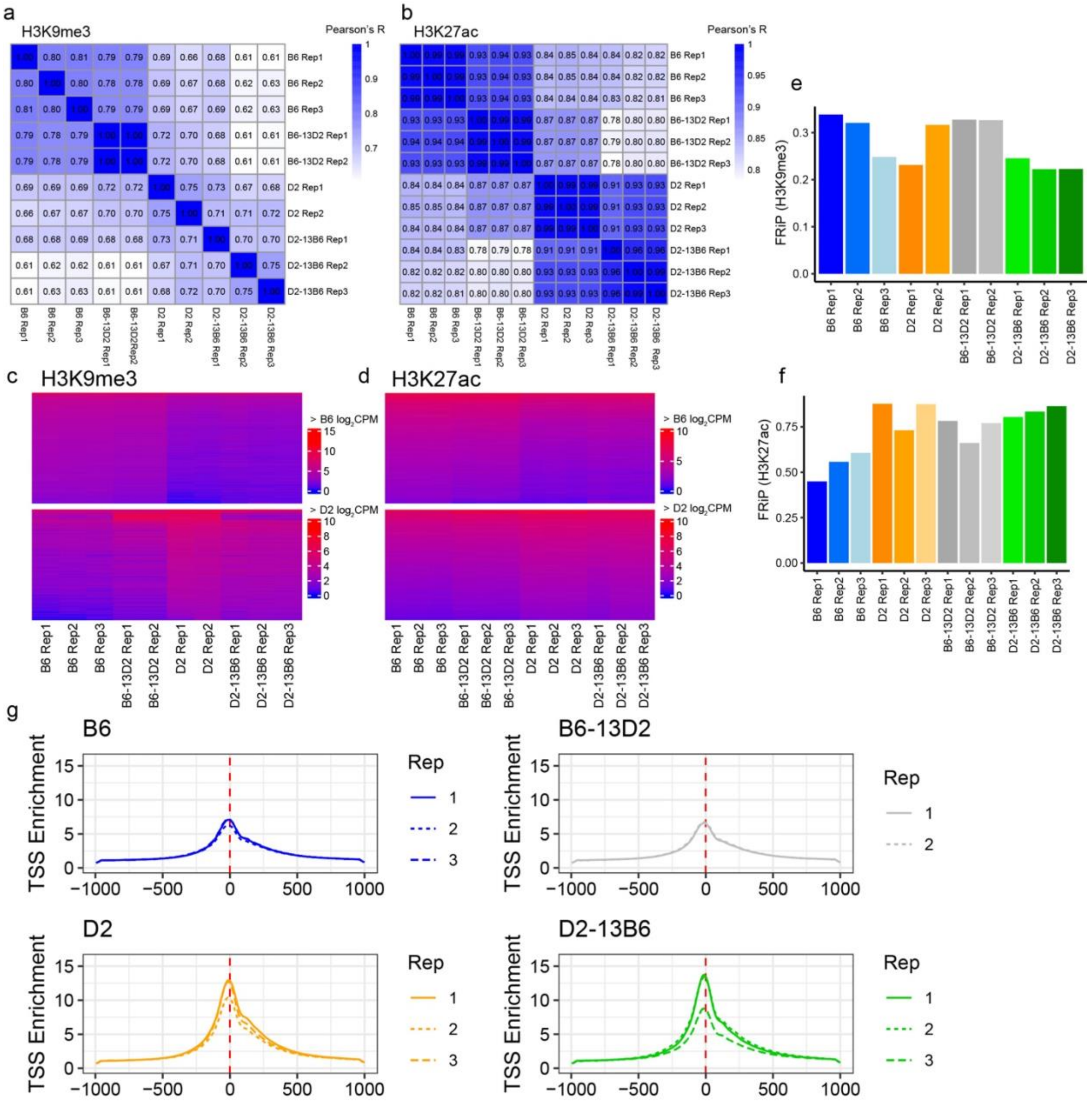


**Fig. S6. Quality control of ChIP-seq and ATAC-seq reveals strong reproducibility among replicates.**
**a** Pearson correlation of H3K27ac and H3K9me3 (**b**) ChIP from technical replicates of mouse embryonic stem cells (ESCs) derived from B6, D2, B6-13D2, and D2-13B6 strains. **c** Heatmap indicating log2CPM read counts from a random sample of 2,500 H3K9me3 or 2,500 H3K27ac (**d**) peaks that are more enriched in B6 (> B6) or more enriched in D2 ( > D2). **e** Fraction of reads in peaks for H3K9me3 and H3K27ac (**f**) ChIP-seq. For H3K9me3, peaks are those identified as ATAC-seq peaks in BXD population. **g** Transcription start site (TSS) enrichment from ATAC-seq data in two or three replicates per strain.

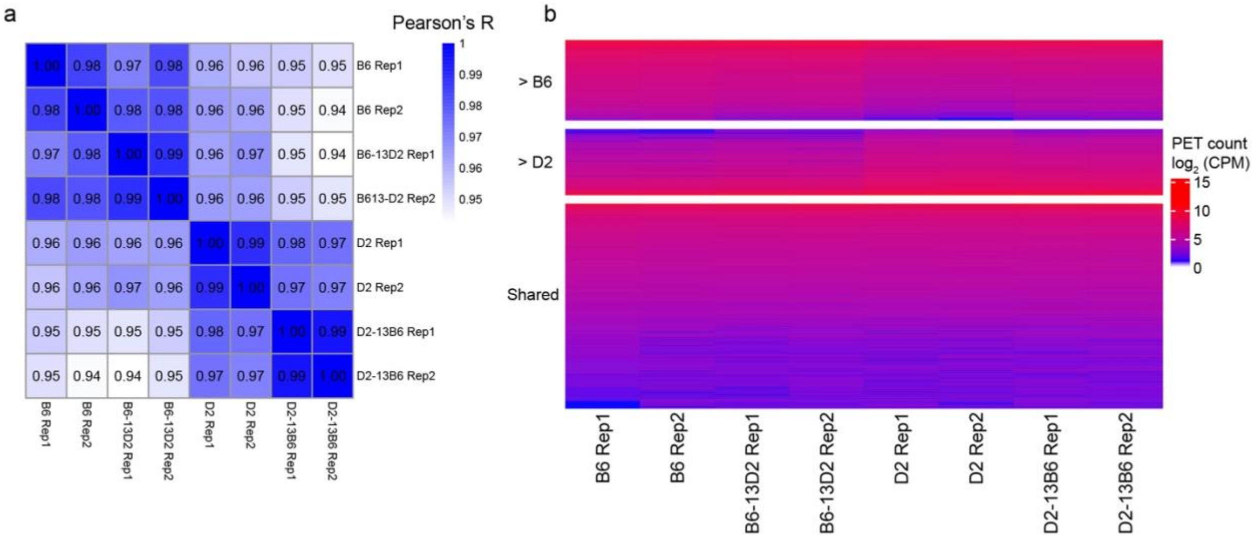


**Fig. S7. Quality control of HiChIP data reveals strong reproducibility among replicates.**
**a** Pearson correlation of H3K27ac HiChIP from technical replicates of mouse embryonic stem cells (ESCs) derived from B6, D2, B6-13D2, and D2-13B6 strains. **b** Heatmap of read counts from 979 interactions that are stronger in B6 (> B6), 806 interactions that are stronger in D2 (> D2) or a random sample of 2,500 interactions that are not significantly different between the two strains (Shared).

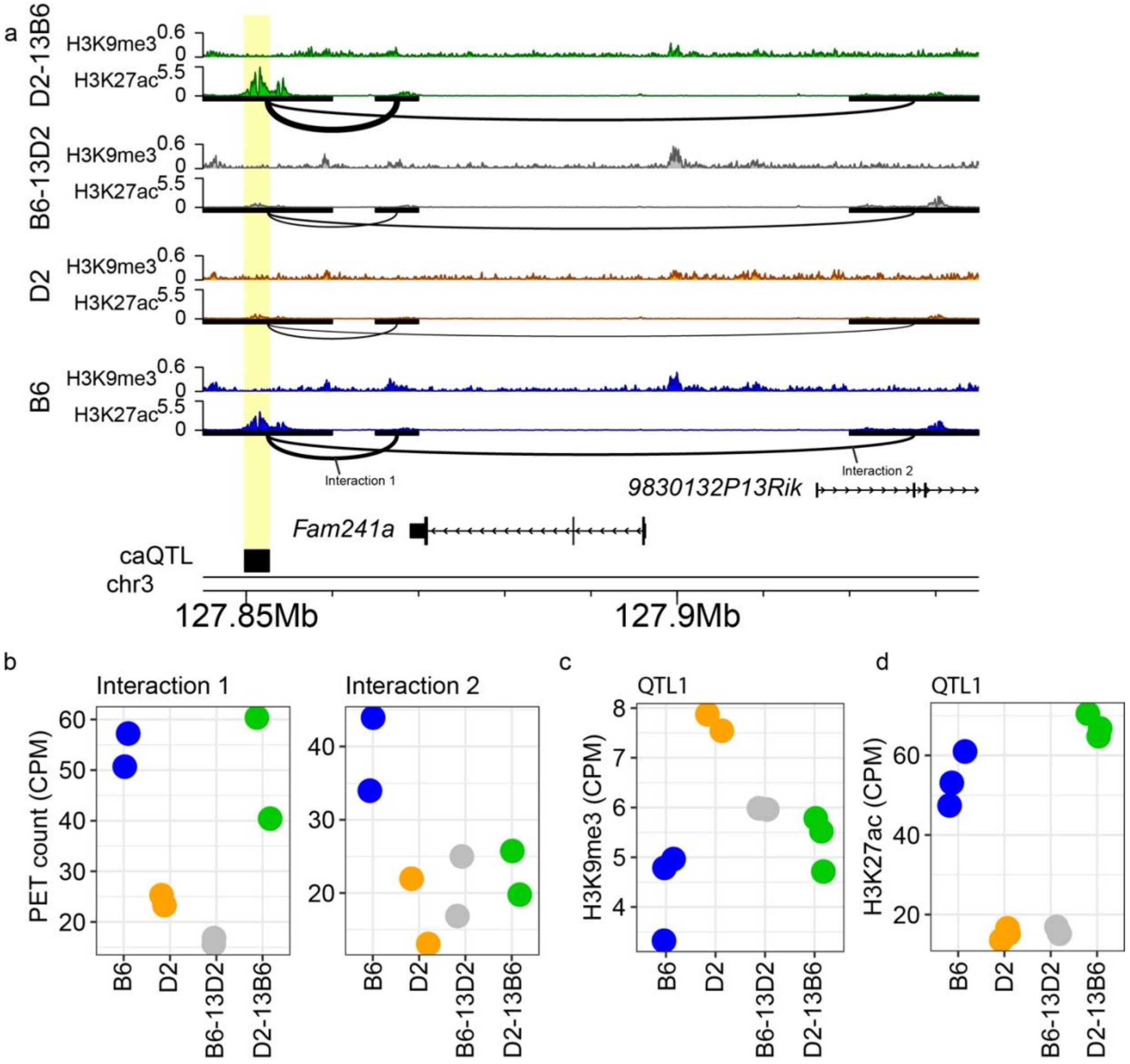


**Fig. S8. Chr13 coordinates heterochromatin and 3D architecture at *Fam241a* locus.**
**a** Genome profiles for H3K27ac and H3K9me3 ChIP enrichment from B6 (blue), D2 (orange), B6-13D2 (grey) and D2-13B6 (green) at the *Fam241a* locus. HiChIP interactions indicated below H3K27ac tracks with the thickness of the arc connecting two anchors representing contact frequency. Highlight indicates a *trans*-target (black box) overlapping a contact anchor and variable histone modifications. **b** HiChIP PET counts for all four strains for two interactions indicated in panel *a* (Interaction 1: BD p = 1.68e-3, BB13D p = 6.48e-7, DD13B p = 1.36e-3; Interaction 2: BD p = 5.73e-3, BB13D p = 1.25e-2, DD13B p = 4.22e-1). **c** H3K9me3 enrichment at Chr13 *trans*-target highlighted panel *a* (BD p = 2.20e-2, BB13D p = 2.19e-1, DD13B p = 1.13e-1). **d** H3K27ac enrichment at Chr13 *trans* caQTL target highlighted panel a (BD p = 5.63e-58, BB13D p = 4.30e-58, DD13B p = 2.66e-80). BD is the comparison between B6 and D2; BB13D is the comparison between B6 and B6-13D2; DD13B is the comparison between D2 and D2-13B6.

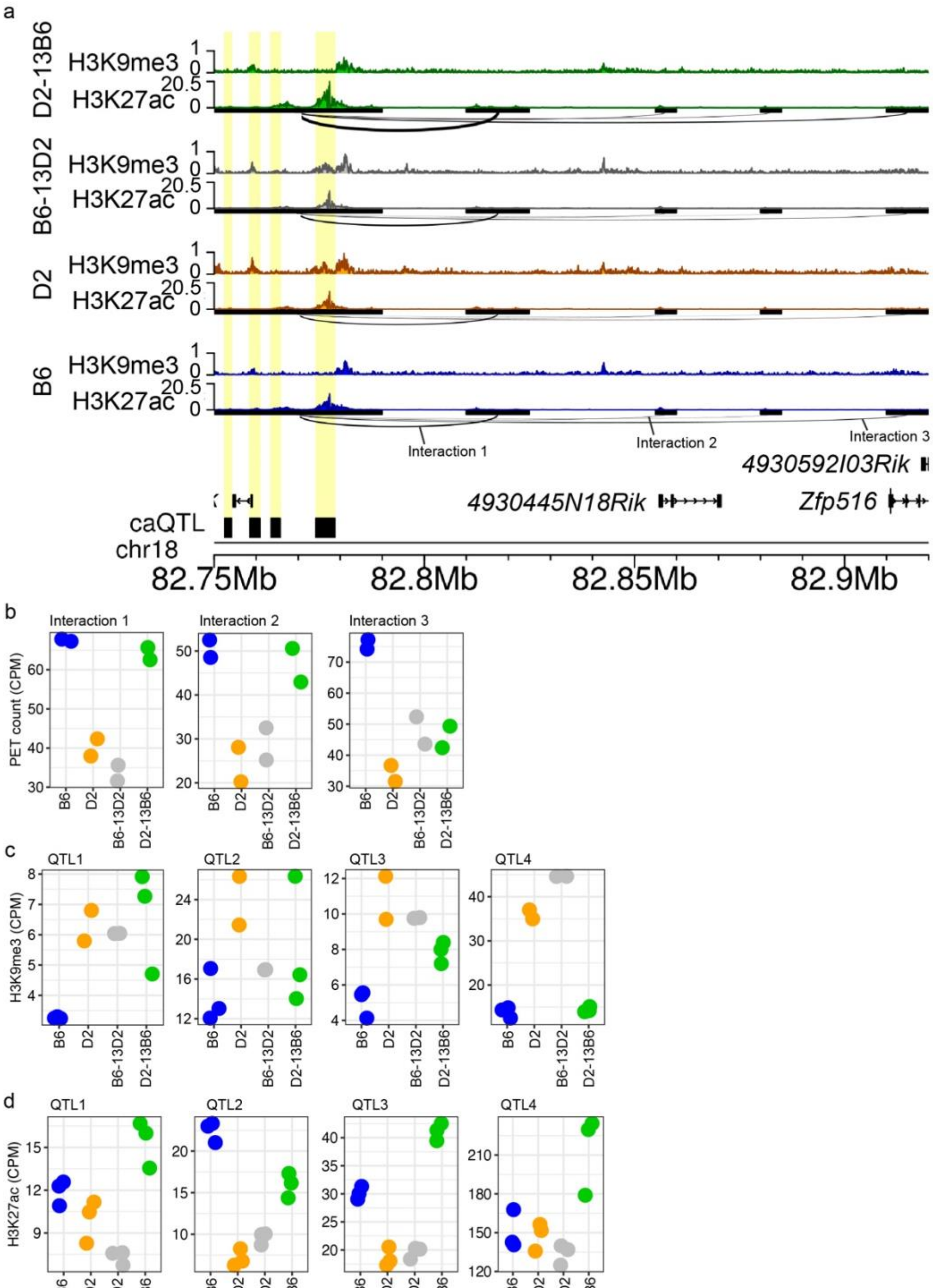


**Fig. S9. Chr13 coordinates heterochromatin and 3D architecture at *Zfp516* locus**.
**a** Genome profiles for H3K27ac and H3K9me3 ChIP enrichment from B6 (blue), D2 (orange), B6-13D2 (grey) and D2-13B6 (green) at the *Zfp516* locus. HiChIP interactions indicated below H3K27ac tracks with the thickness of the arc connecting two anchors representing contact frequency. Highlight indicates a *trans*-target (black box) overlapping a contact anchor and variable histone modifications. **b** HiChIP PET counts for all four strains for two interactions

indicated in panel *a* (Interaction 1: BD p = 1.53e-2, BB13D p = 4.74e-4, DD13B p = 1.60e-2; Interaction 2: BD p = 2.67e-3, BB13D p = 1.08e-2, DD13B p = 3.56e-3; Interaction 3: BD p = 1.79e-4, BB13D p =1.08e-2, DD13B p = 1.75e-1). **c** H3K9me3 enrichment at Chr13 *trans*-targets highlighted panel *a* (QTL1: BD p = 1.69e-2, BB13D p = 2.57e-2, DD13B p = 9.06e-1; QTL2: BD p = 4.91e-3, BB13D p = 3.34e-1, DD13B p = 1.76e-1; QTL3: BD p = 3.02e-4, BB13D p = 1.75e-3, DD13B p = 1.04e-1; QTL4: BD p = 2.08e-13, BB13D p = 1.33e-22, DD13B p = 3.00e-13). **d** H3K27ac enrichment at Chr13 *trans* caQTL target highlighted panel a (QTL1: BD p = 6.96e-2, BB13D p = 4.51e-7, DD13B p = 3.44e-6; QTL2: BD p = 5.14e-35, BB13D p = 2.72e-23, DD13B p = 5.45e-17; QTL3: BD p = 5.14e-35, BB13D p = 2.72e-23, DD13B p = 5.45e-17; QTL4: BD p = 8.24e-1, BB13D p = 7.77e-2, DD13B p = 6.54e-8). BD is the comparison between B6 and D2; BB13D is the comparison between B6 and B6-13D2; DD13B is the comparison between D2 and D2-13B6.

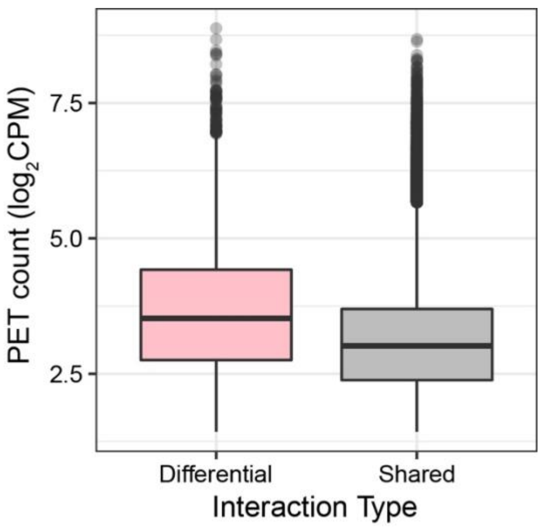


**Fig. S10. Differential interactions show higher PET counts than non-differential interactions.**
**a** Box and whisker plot indicating PET count values from HiChIP for B6 and D2 mESCs for DIs or non-DIs (shared).
