## Supplementary material for "*Trans*-regulation of heterochromatin underlies genetic variation in 3D genome contacts": MiniMUGA Genotyping Results for B6-13D2

### MiniMUGA Background Analysis v2.3.1

**Sample ID**: 13-841

**Neogen ID**: AAAS-5958

**Summary**

The genotype of this sample is of **excellent** quality. It is **male** and **inbred**, and likely a mix of **C57BL/6J** and **DBA/2J**.

Diagnostic SNPs are likely explained by the presence of the background strains

- Solution 1: C57BL/6J and DBA/2J
  - C57BL/6J: 165 / 170 (97.1%)
  - DBA/2J: 3 / 140 (2.1%)

No genetic constructs were detected in this sample.

**Genotyping Quality**

**Excellent (2 N calls)**
All reported results are dependent on genotyping quality.

**Chromosomal Sex**: XY

**Inbreeding Estimate**

100.0% Inbred
(Percentage of the genome (autosomal and X chromosomes) that is homozygous or hemizygous for primary, secondary, and unknown backgrounds. See Genome Analysis)

**Constructs Detected**

| BlastR | bpA | Cas9 | chlor | cHS4 | Cre | DTA | Flp | g_FP | hCMV_a | hCMV_b | hTK_pr | iCre | IRES | Luc | r_FP | rtTA | SV40 | tTA |
|---|---|---|---|---|---|---|---|---|---|---|---|---|---|---|---|---|---|---|
| - | - | - | - | - | - | - | - | - | - | - | - | - | - | - | - | - | - | - |

**Refined Ideogram**

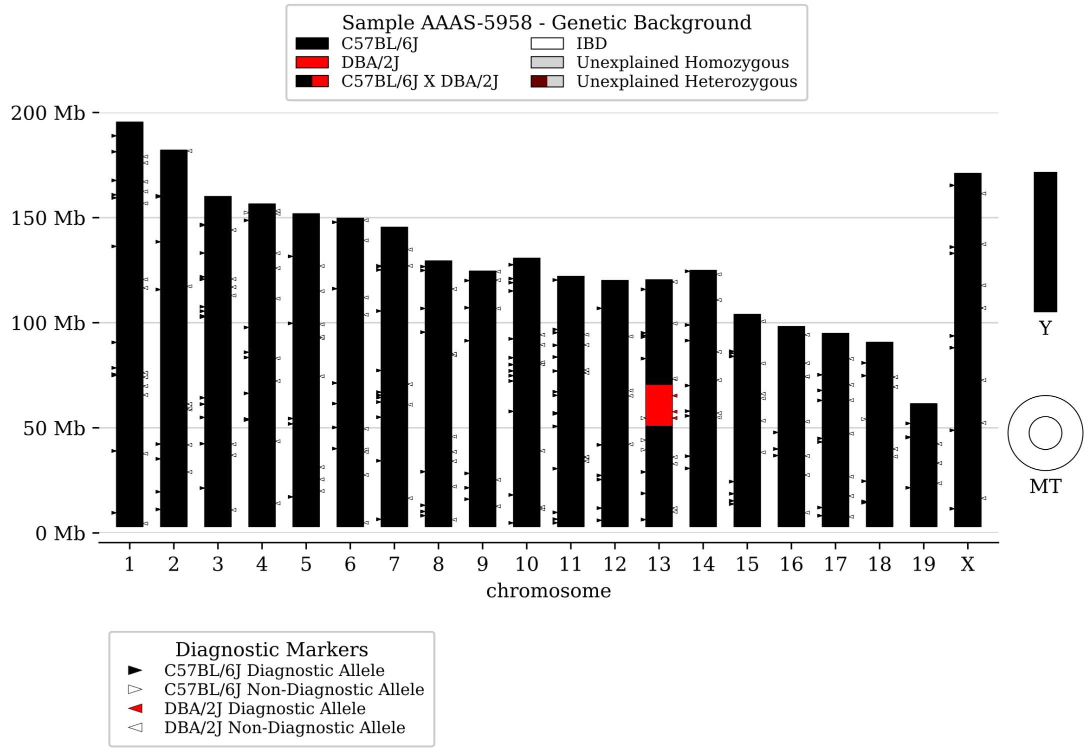


**Genome Analysis**

| Background | Zygosity | Informative Markers | Informative Markers % | Genome % |
|---|---|---|---|---|
| C57BL/6J | Homozygous | 2887 | 99.4% | 99.2% |
| DBA/2J | Homozygous | 16 | 0.6% | 0.8% |
| | **Total** | 2903 | 100.0% | 100.0% |

**Y Chromosome**

Y Haplogroup 11 - 100.0% Consistent
Includes C57BL/6J and 2 other strains

### MiniMUGA Background Analysis v2.3.1

| | |
|---|---|
| **MT Genome** | MT Haplogroup 6 - 100.0% Consistent<br>Includes C57BL/6J, DBA/2J and 166 other strains |

**Backgrounds Detected (Diagnostic Alleles)**

| | Diagnostic Alleles Observed | | | |
|---|---|---|---|---|
| **Diagnostic Class** | **Homozygous** | **Heterozygous** | **Potential** | **% Observed** |
| C57BL/6J, C57BL/6JJicTac, C57BL/6JRj | 101 | 0 | 102 | 99.0% |
| C57BL/6J, C57BL/6JRj | 29 | 1 | 31 | 96.8% |
| C57BL/6J, C57BL/6JEiJ, C57BL/6JJicTac, C57BL/6JRj | 20 | 0 | 21 | 95.2% |
| B6N-Tyr<c-Brd>/BrdCrCrl, C57BL/6J, C57BL/6JJicTac, C57BL/6JRj | 5 | 0 | 5 | 100.0% |
| C57BL/6J | 3 | 0 | 5 | 60.0% |
| B6N-Tyr<c-Brd>/BrdCrCrl, C57BL/6J, C57BL/6JBomTac, C57BL/6JEiJ, C57BL/6JJicTac, C57BL/6JOlaHsd, C57BL/6JRj | 2 | 0 | 2 | 100.0% |
| C57BL/6J, C57BL/6JBomTac, C57BL/6JEiJ, C57BL/6JJicTac, C57BL/6JOlaHsd, C57BL/6JRj | 2 | 0 | 2 | 100.0% |
| DBA/2J | 2 | 0 | 117 | 1.7% |
| B6N-Tyr<c-Brd>/BrdCrCrl, C57BL/6J, C57BL/6JEiJ, C57BL/6JJicTac, C57BL/6JRj | 1 | 0 | 1 | 100.0% |
| C57BL/6J, C57BL/6JEiJ, C57BL/6JJicTac, C57BL/6JOlaHsd, C57BL/6JRj | 1 | 0 | 1 | 100.0% |
| DBA/2J, DBA/2JRj | 1 | 0 | 23 | 4.3% |

**Minimal Strain Sets Explaining All Diagnostic Classes (Number of Markers Explained):**

- Solution 1: C57BL/6J and DBA/2J
    - C57BL/6J: 165 / 170 (97.1%)
    - DBA/2J: 3 / 140 (2.1%)

### MiniMUGA Background Analysis v2.3.1

| | Chromosome | Start (Mb) | Stop (Mb) | Background | Zygosity |
|---|---|---|---|---|---|
| Diplotype Intervals | 1 | 3000000 | 195471971 | C57BL/6J | Homozygous |
| | 2 | 3000000 | 182113224 | C57BL/6J | Homozygous |
| | 3 | 3000000 | 160039680 | C57BL/6J | Homozygous |
| | 4 | 3000000 | 156508116 | C57BL/6J | Homozygous |
| | 5 | 3000000 | 151834684 | C57BL/6J | Homozygous |
| | 6 | 3000000 | 149736546 | C57BL/6J | Homozygous |
| | 7 | 3000000 | 145441459 | C57BL/6J | Homozygous |
| | 8 | 3000000 | 129401213 | C57BL/6J | Homozygous |
| | 9 | 3000000 | 124595110 | C57BL/6J | Homozygous |
| | 10 | 3000000 | 130694993 | C57BL/6J | Homozygous |
| | 11 | 3000000 | 122082543 | C57BL/6J | Homozygous |
| | 12 | 3000000 | 120129022 | C57BL/6J | Homozygous |
| | 13 | 3000000 | 51136860 | C57BL/6J | Homozygous |
| | 13 | 51136860 | 70449416 | DBA/2J | Homozygous |
| | 13 | 70449416 | 120421639 | C57BL/6J | Homozygous |
| | 14 | 3000000 | 124902244 | C57BL/6J | Homozygous |
| | 15 | 3000000 | 104043685 | C57BL/6J | Homozygous |
| | 16 | 3000000 | 98207768 | C57BL/6J | Homozygous |
| | 17 | 3000000 | 94987271 | C57BL/6J | Homozygous |
| | 18 | 3000000 | 90702639 | C57BL/6J | Homozygous |
| | 19 | 3000000 | 61431566 | C57BL/6J | Homozygous |
| | X | 3000000 | 171031299 | C57BL/6J | Hemizygous |
| | Y | 0 | 0 | C57BL/6J | Hemizygous |
| | MT | 0 | 0 | IBD | Hemizygous |
