## Supplementary material for "*Trans*-regulation of heterochromatin underlies genetic variation in 3D genome contacts": MiniMUGA Genotyping Results for D2-13B6

### MiniMUGA Background Analysis v2.3.1

| | |
|---|---|
| **Sample ID** | 14-846 |
| **Neogen ID** | AAAS-5953 |

**Summary**

The genotype of this sample is of **excellent** quality. It is **male** and **inbred**, and likely a mix of **DBA/2J** and (**C57BL/6J and/or C57BL/6JJicTac and/or C57BL/6JEiJ and/or C57BL/6JRj**).

Diagnostic SNPs are likely explained by the presence of the background strains

- Solution 1: C57BL/6J and DBA/2J
  - DBA/2J: 140 / 143 (97.9%)
  - C57BL/6J: 1 / 21 (4.8%)
- Solution 2: C57BL/6JJicTac and DBA/2J
  - DBA/2J: 140 / 143 (97.9%)
  - C57BL/6JJicTac: 1 / 21 (4.8%)
- Solution 3: C57BL/6JEiJ and DBA/2J
  - DBA/2J: 140 / 143 (97.9%)
  - C57BL/6JEiJ: 1 / 21 (4.8%)
- Solution 4: C57BL/6JRj and DBA/2J
  - DBA/2J: 140 / 143 (97.9%)
  - C57BL/6JRj: 1 / 21 (4.8%)

No genetic constructs were detected in this sample.

| | |
|---|---|
| **Genotyping Quality** | **Excellent (6 N calls)**<br>All reported results are dependent on genotyping quality. |
| **Chromosomal Sex** | XY |
| **Inbreeding Estimate** | 100.0% Inbred<br>(Percentage of the genome (autosomal and X chromosomes) that is homozygous or hemizygous for primary, secondary, and unknown backgrounds. See Genome Analysis) |

**Constructs Detected**

| BlastR | bpA | Cas9 | chlor | cHS4 | Cre | DTA | Flp | g_FP | hCMV_a | hCMV_b | hTK_pr | iCre | IRES | Luc | r_FP | rtTA | SV40 | tTA |
|---|---|---|---|---|---|---|---|---|---|---|---|---|---|---|---|---|---|---|
| - | - | - | - | - | - | - | - | - | - | - | - | - | - | - | - | - | - | - |

**Refined Ideogram**

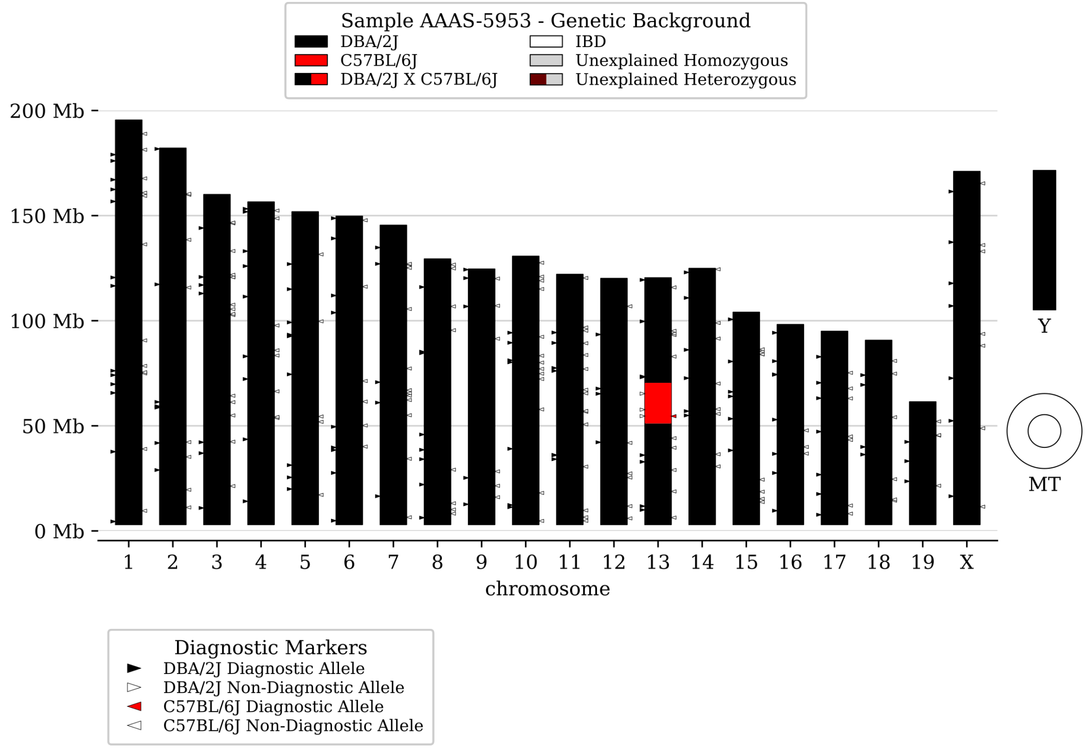

### MiniMUGA Background Analysis v2.3.1

#### Genome Analysis

| Background | Zygosity | Informative Markers | Informative Markers % | Genome % |
|---|---|---|---|---|
| DBA/2J | Homozygous | 2852 | 99.5% | 99.2% |
| C57BL/6J | Homozygous | 15 | 0.5% | 0.8% |
| | **Total** | 2867 | 100.0% | 100.0% |

**Y Chromosome**

Y Haplogroup 9 - 100.0% Consistent
Includes DBA/2J and 21 other strains

**MT Genome**

MT Haplogroup 6 - 100.0% Consistent
Includes C57BL/6J, C57BL/6JEiJ, C57BL/6JJicTac, C57BL/6JRj, DBA/2J and 163 other strains

**Backgrounds Detected (Diagnostic Alleles)**

| | Diagnostic Alleles Observed | | | |
|---|---|---|---|---|
| **Diagnostic Class** | **Homozygous** | **Heterozygous** | **Potential** | **% Observed** |
| DBA/2J | 115 | 0 | 117 | 98.3% |
| DBA/2J, DBA/2JRj | 22 | 0 | 23 | 95.7% |
| DBA/2J, DBA/2JOlaHsd, DBA/2JRj | 3 | 0 | 3 | 100.0% |
| C57BL/6J, C57BL/6JEiJ, C57BL/6JJicTac, C57BL/6JRj | 1 | 0 | 21 | 4.8% |

**Minimal Strain Sets Explaining All Diagnostic Classes (Number of Markers Explained):**

- Solution 1: C57BL/6J and DBA/2J
  - DBA/2J: 140 / 143 (97.9%)
  - C57BL/6J: 1 / 21 (4.8%)
- Solution 2: C57BL/6JJicTac and DBA/2J
  - DBA/2J: 140 / 143 (97.9%)
  - C57BL/6JJicTac: 1 / 21 (4.8%)
- Solution 3: C57BL/6JEiJ and DBA/2J
  - DBA/2J: 140 / 143 (97.9%)
  - C57BL/6JEiJ: 1 / 21 (4.8%)
- Solution 4: C57BL/6JRj and DBA/2J
  - DBA/2J: 140 / 143 (97.9%)
  - C57BL/6JRj: 1 / 21 (4.8%)

### MiniMUGA Background Analysis v2.3.1

| | Chromosome | Start (Mb) | Stop (Mb) | Background | Zygosity |
|---|---|---|---|---|---|
| **Diplotype Intervals** | 1 | 3000000 | 195471971 | DBA/2J | Homozygous |
| | 2 | 3000000 | 182113224 | DBA/2J | Homozygous |
| | 3 | 3000000 | 160039680 | DBA/2J | Homozygous |
| | 4 | 3000000 | 156508116 | DBA/2J | Homozygous |
| | 5 | 3000000 | 151834684 | DBA/2J | Homozygous |
| | 6 | 3000000 | 149736546 | DBA/2J | Homozygous |
| | 7 | 3000000 | 145441459 | DBA/2J | Homozygous |
| | 8 | 3000000 | 129401213 | DBA/2J | Homozygous |
| | 9 | 3000000 | 124595110 | DBA/2J | Homozygous |
| | 10 | 3000000 | 130694993 | DBA/2J | Homozygous |
| | 11 | 3000000 | 122082543 | DBA/2J | Homozygous |
| | 12 | 3000000 | 120129022 | DBA/2J | Homozygous |
| | 13 | 3000000 | 51136860 | DBA/2J | Homozygous |
| | 13 | 51136860 | 70449416 | C57BL/6J | Homozygous |
| | 13 | 70449416 | 120421639 | DBA/2J | Homozygous |
| | 14 | 3000000 | 124902244 | DBA/2J | Homozygous |
| | 15 | 3000000 | 104043685 | DBA/2J | Homozygous |
| | 16 | 3000000 | 98207768 | DBA/2J | Homozygous |
| | 17 | 3000000 | 94987271 | DBA/2J | Homozygous |
| | 18 | 3000000 | 90702639 | DBA/2J | Homozygous |
| | 19 | 3000000 | 61431566 | DBA/2J | Homozygous |
| | X | 3000000 | 171031299 | DBA/2J | Hemizygous |
| | Y | 0 | 0 | DBA/2J | Hemizygous |
| | MT | 0 | 0 | IBD | Hemizygous |
